# Targeting FOXA1 and FOXA2 Disrupts the Lineage-Specific Oncogenic Output Program in Prostate Cancer

**DOI:** 10.1101/2025.08.21.671500

**Authors:** Nicolo Formaggio, Jacopo Sgrignani, Gayathri Thillaiyampalam, Claudio Lorenzi, Giada A. Cassanmagnago, Marco Coazzoli, Federico Costanzo, Yanick Uebelhart, Daniela Bossi, Diego Camuzi Cassiano, Andrea Rinaldi, Matteo Pecoraro, Roger Geiger, Raffaella Santoro, Marco Bolis, Andrea Cavalli, Jean-Philippe Theurillat

## Abstract

Activation of the androgen receptor (AR) is the key lineage-specific oncogenic pathway and the primary therapeutic target in prostate cancer. While AR signaling is enabled by the pioneer transcription factor FOXA1, its homolog FOXA2 is specifically expressed in advanced lineage plasticity prostate cancers that have lost the AR signaling axis. However, their roles and utility as drug targets remain incompletely characterized. Here, we show an unexpected collaboration of FOXA1 and FOXA2 in mediating AR-independent cell proliferation in different lineage plasticity cancer subtypes. Conversely, joint loss-of-function or pharmacologic disruption of FOXA1 and FOXA2 leads to the collapse of lineage-specific oncogenic transcription factors followed by cell cycle arrest. In summary, our findings uncover a druggable dependency for AR-positive and - negative prostate cancers.

## INTRODUCTION

The androgen receptor (AR) is a critical oncogenic lineage-specific pathway in prostate cancer ^1,2^. Upon binding to androgens, AR dimerizes and enters the cell nucleus, activating a transcriptional output program that supports tumor cell proliferation ^3^. Consequently, androgen deprivation therapy (ADT) and more recent androgen receptor signaling inhibitors (ARSi) (e.g., second-generation anti-androgens such as enzalutamide or the androgen synthesis inhibitor abiraterone) remain the cornerstone of prostate cancer therapy in an early metastatic setting ^2,4,5^. However, the harsher inhibition of AR by ARSi has led to an increase in prostate cancers that lose this pathway through lineage plasticity ^6,7^. The latter includes a variety of new emerging subtypes of AR-negative prostate cancers, such as neuroendocrine prostate cancer (NEPC) and double-negative prostate cancers (DNPC) that are characterized by either activation of WNT-signaling or a stem-cell-like (SCL) phenotype^7–9^.

FOXA1 is an established pioneer transcription factor for steroid hormone receptors and a key selective dependency in prostate and breast cancer (https://depmap.org) ^10–16^. In prostate cancer, its homolog FOXA2 is specifically upregulated in advanced lineage-plasticity prostate cancers that have lost the androgen receptor (AR) signaling axis ^17–22^. Nevertheless, FOXA1 and FOXA2 functions have not been assessed in the very same cellular context thus far. In this study, we characterize the chromatin occupancy of FOXA1 and FOXA2 in various prostate cancer models and uncover a surprising collaboration for FOXA2 and FOXA1 in promoting AR independence, which can be targeted by small compounds capable of destabilizing the forkhead domains of FOXA1/2 in concert, and thereby, disrupting various lineage-specific oncogenic pathways in this disease.

## RESULTS

### FOXA1 and FOXA2 co-localize at the chromatin in lineage-plasticity prostate cancers

To address the expression pattern of FOXA1 and FOXA2 in advanced prostate cancer, we consulted a recently harmonized bulk and single-cell RNA sequencing (RNAseq) atlas ^23^ (www.prostatecanceratlas.org). While FOXA2 was exclusively expressed in NEPC and DNPC, we noticed, in line with a previous study, that most tumors also co-expressed FOXA1, however, at a somewhat lower level than in AR-positive castration-resistant prostate cancers (ARPC) (Figures 1A & S1A, B) ^24^. In agreement, FOXA1 and FOXA2 co-expression was found in different cancer cell clusters of NEPC and WNT-driven and SCL-subtypes of DNPC, as reported recently (Figures S1C-E) ^19^. We observed a similar pattern of FOXA1 and FOXA2 protein abundance in different models of lineage plasticity cancers representing SCL (PC3), WNT-driven prostate cancers (MSK-PCa1), and NEPC (H660, LuCaP-145.2) as compared to AR-driven LNCaP, LAPC4 and 22Rv1 cells (Figure 1B).

**Figure 1.**
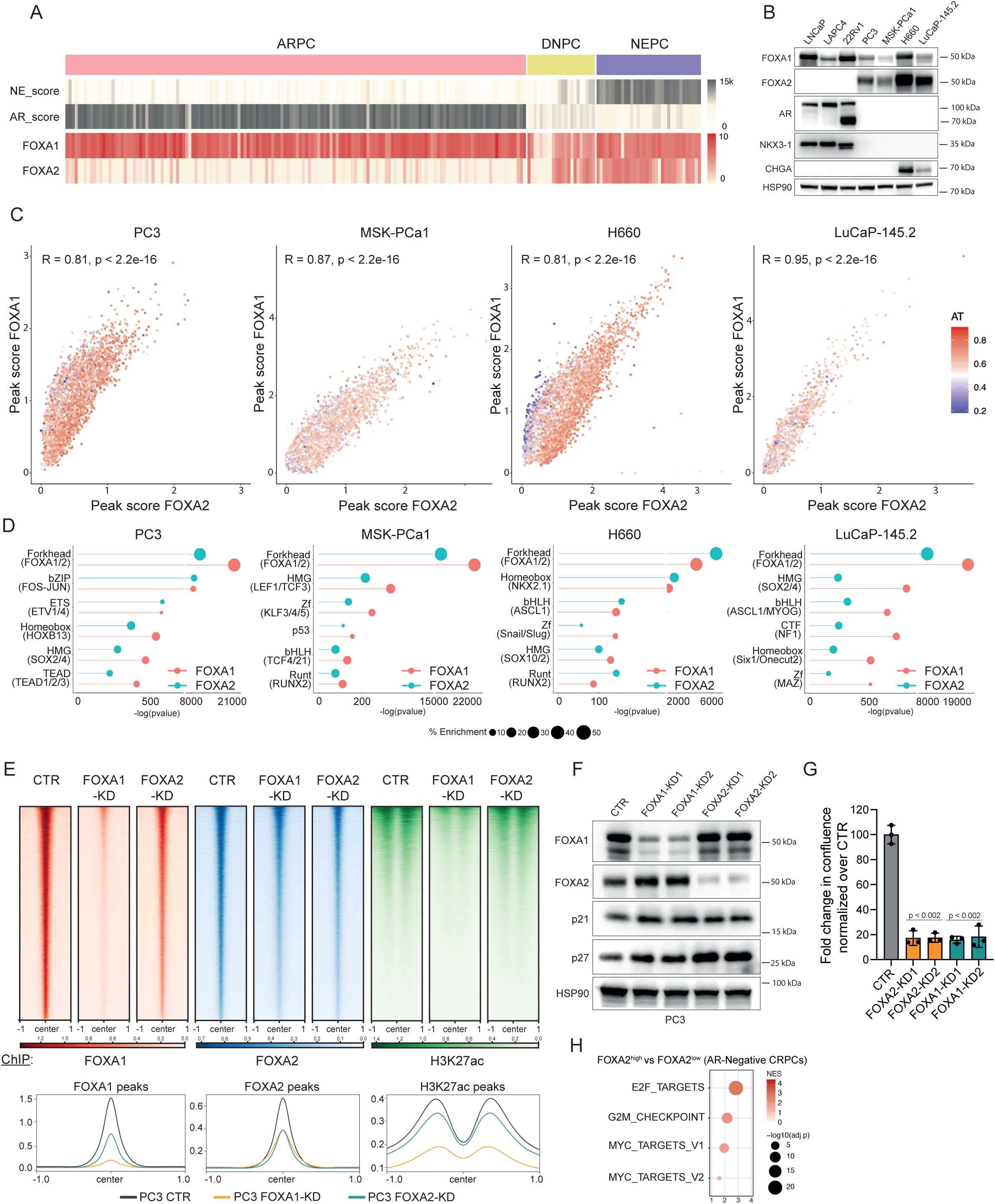
FOXA1 and FOXA2 colocalize at the chromatin in lineage plasticity prostate cancers. **(A)** Bulk RNA sequencing atlas data highlights FOXA1 and FOXA2 expression levels in AR-positive castration-resistant prostate cancer (ARPC), neuroendocrine prostate cancer (NEPC), and double-negative prostate cancer (DNPC) (www.prostatecanceratlas.org^23^). Androgen receptor (AR) and neuroendocrine (NE) scores are presented as previously published ^7^. **(B)** Immunoblot for the indicated proteins, in the indicated cell lines (LNCaP, LAPC4, 22Rv1, H660), and patients-derived xenograft (LuCaP-145.2) and organoid (MSK-PCa1) models. **(C)** Scatter plots showing peak scores for FOXA1 and FOXA2 determined by ChIPseq in the indicated models mentioned above using one replicate each. Pearson correlation coefficient and corresponding p-values are indicated. The AT-content of the peaks are color-coded as indicated on the right side. **(D)** Corresponding motif enrichment analysis (see also table S1_1) highlighting relevant enriched DNA motifs (upper part) and exemplified corresponding transcription factors (in brackets below). **(E)** ChIPseq heatmaps and corresponding global profiles for FOXA1, FOXA2, and H3K27ac of control PC3 cells and knockdown (KD) for FOXA1 or FOXA2 using one replicate each (see Figure S4A for second replicate). **(F)** Immunoblot for the indicated proteins of PC3 control cells and knockdown for FOXA1 and FOXA2 using two independent hairpin RNAs (KD1, KD2).**(G)** Changes in cell proliferation measured by Incucyte® upon knockdown of FOXA1 or FOXA2 as indicated. P-values <0.002 are referred to each KD over CTR (student’s t-test). **(H)** Gene set enrichment analysis of AR-negative clinical prostate cancer samples stratified into FOXA2 high and low expressing cancers (see also Figure 1A & table S1_2). The FDR q-values indicate the significance of the indicated gene sets.

To elucidate the role of FOXA1 and FOXA2 in lineage plasticity prostate cancers, we determined the chromatin occupancy of FOXA1 and FOXA2 by chromatin immunoprecipitation followed by sequencing (ChIPseq) in the different models. We consistently observed a striking co-localization of FOXA1 and FOXA2 at the chromatin (Figure 1C). The FOXA1/2 peaks were enriched for motifs of oncogenic transcription factors related to the different subtypes of prostate cancer (e.g., AP1 and HOXB13 in SCL, NKX2-1 and ASCL1 in NEPC, TCF/LEF in WNT) and were widely distributed across the genome (Figures 1D & S1F, G; table S1_1).

To investigate if FOXA1 and FOXA2 collaborate in oncogenesis, we assessed in PC3 cells if either FOXA1 or FOXA2 knockdown impaired not only the chromatin binding of the corresponding homolog but also H3K27 acetylation (H3K27ac), a marker of chromatin opening. Indeed, FOXA1 knockdown substantially reduced FOXA2 chromatin binding and vice versa (Figures 1E, F). In each case, the reduction of FOXA1 or FOXA2 was paralleled with a decrease in H3K27ac, suggesting that both pioneer factors contribute jointly to chromatin opening and oncogenic transcription. In agreement, the knockdown of either of the factors reduced cell proliferation and led to cell cycle block, as evidenced by an increase in p21 and p27 expression (Figures 1F, G & S1H). FOXA1 has been shown previously to open chromatin and promote survival in H660 cells ^24^. Complementing data for FOXA2 aligned with the findings in PC3 cells (Figure S2). Finally, mining our human bulk RNA data revealed increased cell proliferation as a hallmark of lineage plasticity in prostate cancers expressing FOXA2 (Figures 1H & table S1_2).

### FOXA1 and FOXA2 synergize in driving AR independence

To further explore the interplay of FOXA1 and FOXA2 related to tumorigenesis, we chose to over-express FOXA2 in otherwise negative LNCaP cells and assess for AR-independent growth using either enzalutamide (ENZA) or the AR degrader ARCC4. Both perturbations significantly decreased the levels of the AR target NKX3-1 while ARCC4 strongly reduced AR protein abundance, as expected (Figure 2A). Importantly, forced expression of FOXA2 strongly promoted the growth of LNCaP cells under AR suppression without affecting FOXA1 protein abundance (Figures 2A, B & S3A). Moreover, prolonged pharmacologic degradation of AR increased FOXA2 protein abundance and chromatin occupancy in the resistant cell lines (Figures S3B, C). A similar but slightly lower increase in FOXA2 protein abundance was also noticed upon longer-term ENZA exposure (Figure S3D). In both cases, changes at the protein levels were not related to gene expression (Figure S3E), suggesting a possible increase in protein stability due to enhanced chromatin engagement. Next, we performed RNAseq to study possible pathways triggered by FOXA2 under androgen-rich medium conditions or under the suppression of AR by ENZA or ARCC4. In each case, gene sets related to the cell cycle were among the most enriched, notably when the AR response gene set was reduced (Figures 2C & S3F, table S2). The data aligns well with the previous human data analysis (Figure 1H).

**Figure 2.**
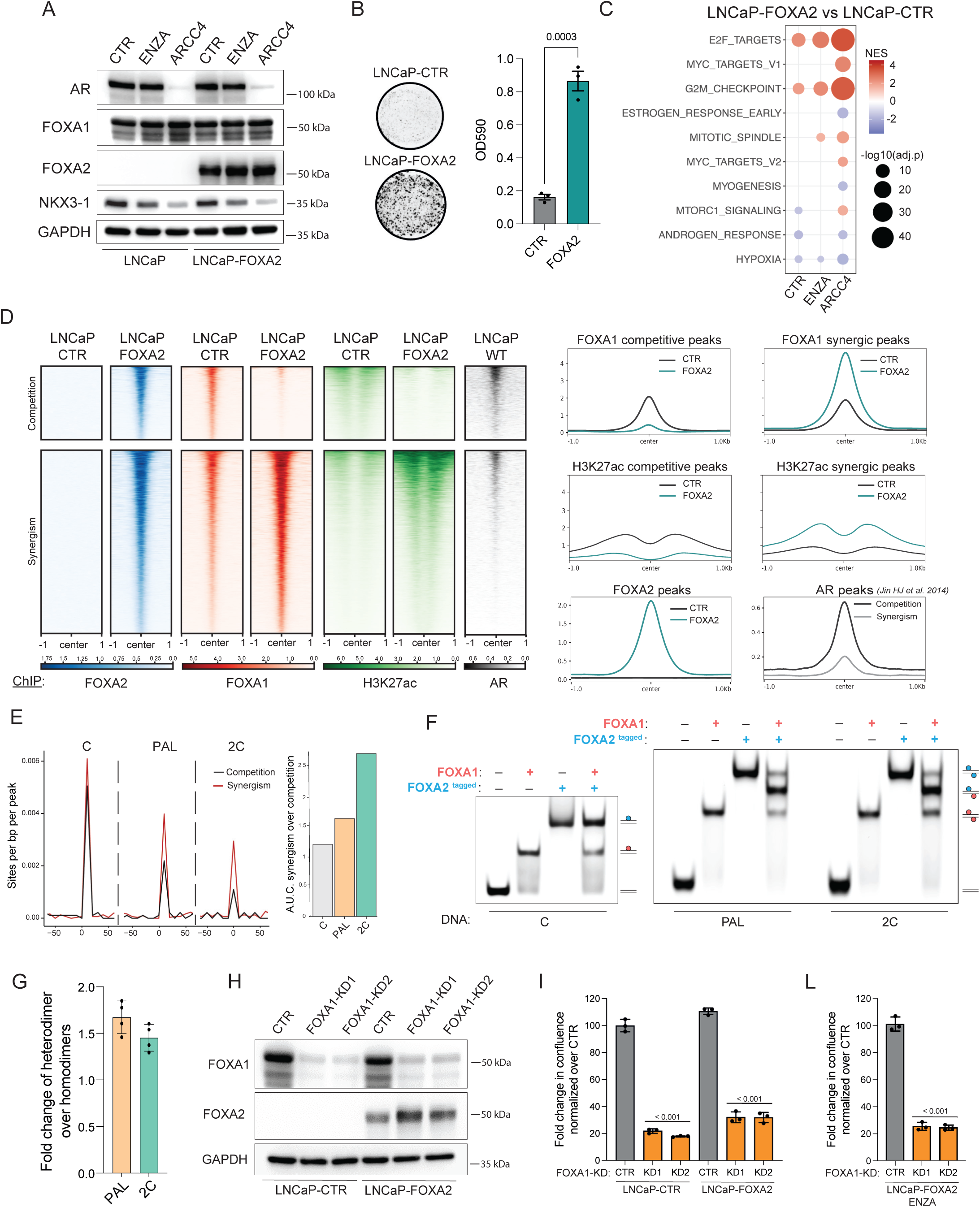
FOXA2 over-expression in LNCaP cells promotes AR-independent growth. **(A)** Immunoblot for the indicated proteins of LNCaP control cells and cells over-expressing FOXA2 treated with DMSO control, enzalutamide (ENZA) 10µM or the AR degrader ARCC4 1µM. **(B)** Corresponding clonogenic assay of LNCaP control (CTR) and FOXA2-over-expressing cells (LNCaP-FOXA2) under ARCC4 1µM treatment. P-value 0.001 of FOXA2 over CTR (student’s t-test). **(C)** Corresponding gene set enrichment analysis of FOXA2 over-expressing cells compared to control cells under DMSO, ENZA 10µM, or ARCC4 1µM treatment. The FDR q-values indicate the significance of the indicated gene sets. **(D)** ChIPseq heatmaps and global profiles of LNCaP control cells and over-expressing FOXA2 for FOXA1, FOXA2, and H3K27ac. AR ChIPseq data of parental LNCaP cells was culled from the literature ^25^. Competition refers to peaks of FOXA2 which decrease for FOXA1 and H3K27ac, while synergism refers to FOXA2 peaks where FOXA1 and H3K27ac are increasing (see also Figure S4E for further stratification). **(E)** Prevalence of FOXA-motifs in chromatin areas of FOXA2-induced competition and synergism in LNCaP cells (left). Changes in the area under the curve (A.U.C) are quantified on the right. **(F)** Electrophoretic Mobility Shift Assay (EMSA) using the canonical FOXA DNA motif (C), the extended palindromic DNA motif (PAL), and the two canonical DNA motifs separated by three nucleotides (2C) as DNA inputs (100nM) and untagged FOXA1 (400nM) and Twin-Strep-tagged FOXA2 (400nM) forkhead domains. When used together, the concentration of both proteins was reduced to 200nM. See also Figure S5A. **(G)** Quantification of the homodimers versus heterodimers (e.g., upper and lower band versus middle band) for PAL and 2C DNA oligonucleotides as indicated. **(H)** Immunoblot for the indicated proteins of LNCaP control cells and cells overexpressing FOXA2 in the presence and absence of FOXA1 knockdown (KD) using two hairpin RNA for each gene as indicated. **(I)** The corresponding change in cell proliferation was measured by Incucyte® upon knockdown of FOXA1 and/or over-expression of FOXA2, as indicated. P-values <0.001 for each KD over their CTR (student’s t-test). **(J)** Change in the cell proliferation measure by Incucyte® in FOXA2 over-expressing LNCaP (LNCaP-FOXA2) treated with 10µM ENZA upon knockdown of FOXA1 using two hairpins as indicated. P-values<0.001 for each KD over CTR (student’s t-test).

To elucidate related changes at the chromatin level, we performed ChIPseq for FOXA1, FOXA2, and H3K27ac in LNCaP cells with previously annotated AR binding sites ^25^. On this occasion, we validated and secured the specificity of both FOXA1 and FOXA2 antibodies. FOXA2 peaks were only detected upon FOXA2 over-expression in LNCaP cells (Figure 2D). Conversely, FOXA1 peaks were not detected in a human fibroblast line (HS-5) over-expressing FOXA2 (Figure S3G-I).

Next, we determined chromatin changes when FOXA2 is over-expressed in LNCaP cells. Interestingly, we noticed a reduction of FOXA1 occupancy and H3K27ac at sites enriched for AR in a previously published dataset (referred to as competition, Figure 2D). In contrast, FOXA1 and FOXA2 strongly co-localized in the bulk part of the genome where they paralleled with increased H3K27ac (referred as synergism) in agreement with the previous data of lineage plasticity models (Figures 1, 2D & S4B-D). The few FOXA1 peaks with no changes did not show any correlation to previously published AR sites and were mainly localized in areas of low H3K27ac (Figure S4E).

The FOXA1 forkhead domain has been shown to bind in different configurations to DNA sequences. Apart from binding to a canonical motif (C), the forkhead domains can also homo-dimerize by binding on the opposite strands of either an extended palindromic motif (PAL) or on two canonical motifs separated by three nucleotides (2C) (Figure S5A)^26^. Thus, we wondered if the synergistic peaks of FOXA1/2 upon FOXA2 over-expression are enriched in the latter motifs. Indeed, interrogating the ChIPseq data revealed that both divergent and convergent motifs were enriched in these regions compared to loci that underwent competition (Figure 2E).

We analyzed whether the FOXA2 forkhead domain can also form homodimers and, more importantly, heterodimers with FOXA1. To this end, FOXA2 showed a similar concentration-dependent engagement with the above-mentioned three oligonucleotide sequences (Figures S5B, C). To test the possibility of heterodimeric complexes between FOXA1 and FOXA2, we used an untagged FOXA1 forkhead domain and a corresponding tagged (Twin-Strep) version of FOXA2 that can be differentiated by molecular size and repeated the EMSA assay (Figure 2F). While results demonstrate that both forkhead domains can homo- and heterodimerize, the quantification of the bands corresponding to homo-versus heterodimers across multiple replicates revealed a preference for the latter. In aggregate, the data suggests that FOXA2 over-expression relocates FOXA1 to areas of the chromatin where they form increasingly heterodimers.

We finally assessed whether FOXA1 becomes dispensable when FOXA2 is over-expressed in LNCaP cells. In opposition to a redundant function, FOXA1 knockdown severely affected the growth of LNCaP cells regardless of FOXA2 in either androgen-rich conditions or in the presence of ENZA (Figures 2G-I). In aggregate, the data further supports the collaboration of FOXA1 and FOXA2 in driving AR-independent growth.

### FOXA1 and FOXA2 maintain the core components of the AR transcriptional machinery

Next, we mapped the proteins in the vicinity of FOXA1 and FOXA2 using proximity-dependent Biotin Identification (BioID) PC3, H660, and LNCaP over-expressing FOXA2 ^27^. For this purpose, we over-expressed either a doxycycline-inducible control nuclear GFP or FOXA1/2 fused to biotin ligase and secured nuclear localization for FOXA1/2 and protein biotinylation in streptavidin pull-down assays as shown for PC3 cells (Figures S6A-E). Overall, the emerging data showed a robust agreement of the absolute values of enriched proteins emerging from GFP control, FOXA1 and FOXA2 BioID in all cell lines (Figure S7A). Moreover, chromatin-related processes were enriched in both FOXA1 and FOXA2 BioID compared to the GFP control, as expected (Figure S7B). Moreover, we noticed a strong overlap of enriched proteins of FOXA1 and FOXA2 in each cellular setting (Figure 3A & table S3_1). The presence of FOXA1 in FOXA2 BioID and vice versa was also confirmed by biotin pulldown, followed by immunoblotting in PC3 cells (Figure S7C). Finally, the FOXA1 BioID results in LNCaP cells were largely superimposable in LNCaP-FOXA2 compared to LNCaP-CTR cells (Figure S7D).

**Figure 3.**
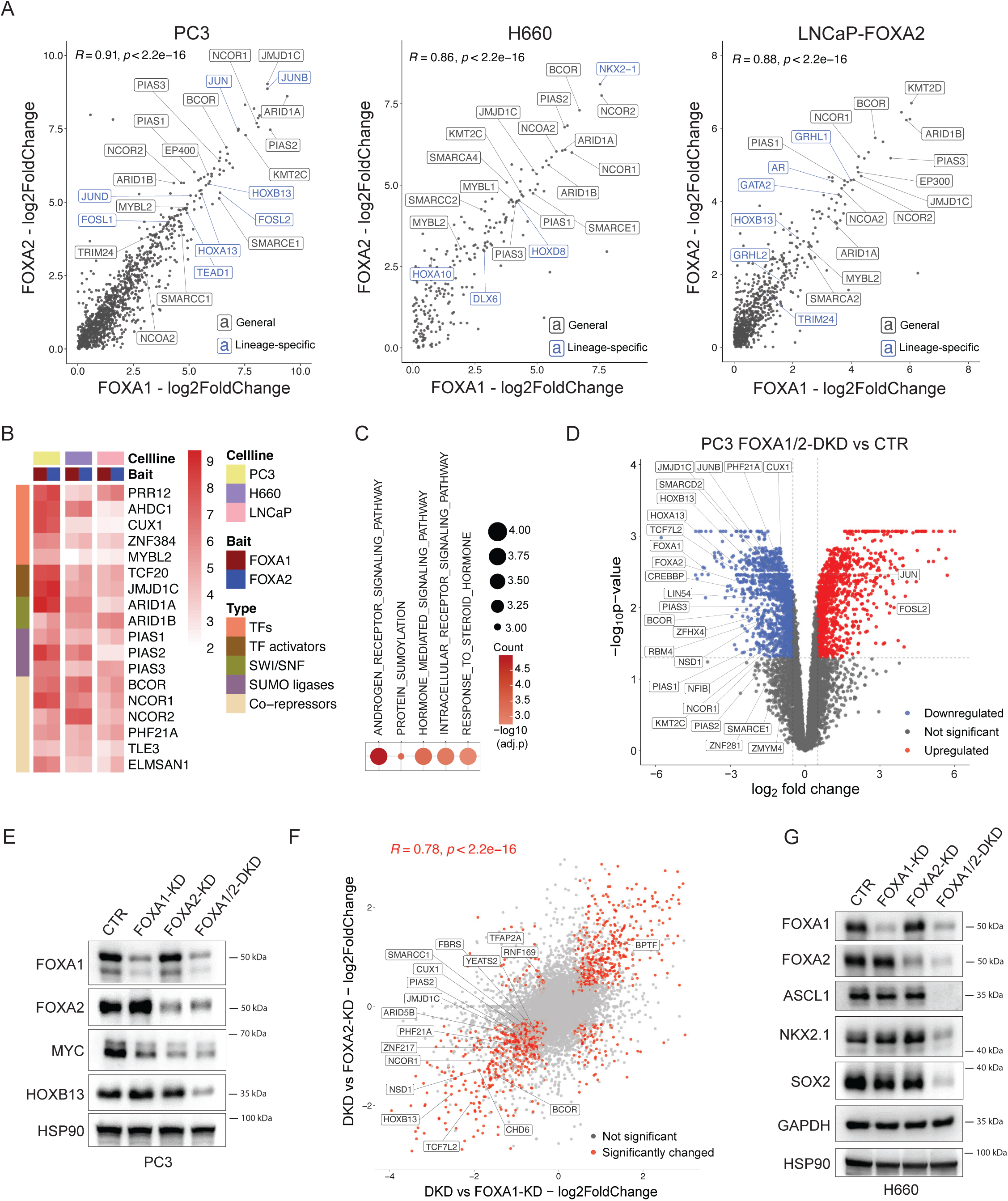
Proximity-dependent Biotin Identification (BioID) interactors of FOXA1 and FOXA2 are largely superimposable and share core components related to AR signaling. **(A)** Scatter plots for the indicated cell lines comparing enrichment scores for FOXA1 and FOXA2 BioID over GFP control. Lineage-specific transcription factors and general core components are specifically labelled as indicated among other proteins. Pearson correlation coefficient and corresponding p-values are indicated. See also Figure S7A for raw values. **(B)** Heatmap of the log2 enrichment scores of the shared eighteen proteins derived from the top 100 enriched common proteins across PC3, H660 and LNCaP-FOXA2 cells in the indicated cell lines and indicated baits. The function of the individual proteins is annotated by color code on the left. **(C)** Corresponding enrichment scores of gene ontology signatures derived from the common 18 proteins between PC3, H660 and LNCaP-FOXA2. The FDR q-values indicate the significance of the indicated gene sets. **(D)** Volcano plot of up and down-regulated proteins comparing FOXA1/2 double knockdown to control using mass-spectrometry. Selected proteins are highlighted (see also Figure S8C-E and table S3_3). **(E)** Immunoblot analysis of the indicated proteins of PC3 with single and double knockdown (KD) of FOXA1/2 using one hairpin RNA each. **(F)** Corresponding scatter plot comparing FOXA1/2 double knockdown to single knockdown. Selected top 100 BioID proteins are highlighted (see table S3_3). Pearson correlation coefficient and corresponding p-values are indicated. **(G)** Corresponding immunoblot analysis of H660 cells for the indicated proteins with single and double knockdown (KD) for FOXA1/2.

In each cell line, we observed the enrichment of key transcription factors characterized for each prostate cancer subtype, i.e., AP1 complex members and HOXB13 in PC3 cells (SCL), NKX2-1in H660 cells (NEPC), and AR, HOXB13 in LNCaP cells (ARPC) ^9,10,16,24,28,29^. We noted a significant overlap of proteins shared across all three cell types for FOXA1 and FOXA2 among the top 100 enriched proteins (Figures S8A, B). Among these BioID core components, most were shared in FOXA1 and FOXA2 BioID (Figure 3B). Importantly, gene ontology analysis revealed a strong link with AR and more broadly steroid hormone signaling, suggesting that even in the absence of AR, FOXA1 and FOXA2 maintain core components of the AR transcriptional machinery throughout disease progression (Figures 3C, S8C, D & table S3_2).

We wondered if proteins in the proximity of FOXA1 and FOXA2 may change abundance with the knockdown of FOXA1 and/or FOXA2 and tested this in PC3 cells. While the single knockdown only slightly affected the protein abundance of the top 100 BioID proteins by mass-spectrometry, double-knockdown significantly reduced the latter (Figures 3D-F, S8E-G & table S3_3). Matched RNAseq data revealed that the reduction of BioID protein abundance could only be partially explained by corresponding gene expression changes (Figures S8H, I & table S3_3). Importantly, we found a similar reduction pattern of the abundance of lineage-specific transcription factors upon double-knockdown of FOXA1/2 in H660 cells (e.g., NKX2-1, SOX2, ASCL1) (Figure 3G). In agreement, the double-knockdown of FOXA1 and FOXA2 reduced cell proliferation more significantly than the single knockdown for one or the other protein in both cell lines (Figures S9A, B). Finally, we didn’t find evidence for upregulation of AR protein abundance or downstream targets (e.g., NKX3-1) upon single or double-knockdown in PC3 and H660 cells (Figures S9C, D).

### The forkhead domains of FOXA1 and FOXA2 can be disrupted by small molecules

Given the similarity of the forkhead domains of FOXA1 and FOXA2 and their DNA binding motifs, we set out to search for small molecules that disrupt both transcription factors jointly. For this purpose, we performed a virtual drug screen using a diversity-driven 54k library on an area of FOXA2 with multiple surface pockets or grooves adjacent to where the DNA binds (Figures 4A & S10A) ^30^. The emerging top nineteen compounds were subsequently tested for their ability to reduce the activity of a FOXA-driven luciferase reporter assay (Figures S10B, C) ^12^. Out of these, three compounds reduced at 100µM the luciferase activity induced by either FOXA1 or FOXA2 by more than 50 % (referred to as S, T, and U) (Figures 4B & S10C). The T compound also reduced the luciferase reporter assay driven by the homolog FOXA3 (Figure S10D), indicating that the current compound T is not specific to FOXA1 and FOXA2. While FOXA3 is clearly less expressed in human prostate cancer tissues (Figure S10E), the data demonstrate potential for improvement of the specificity of the current compounds.

**Figure 4.**
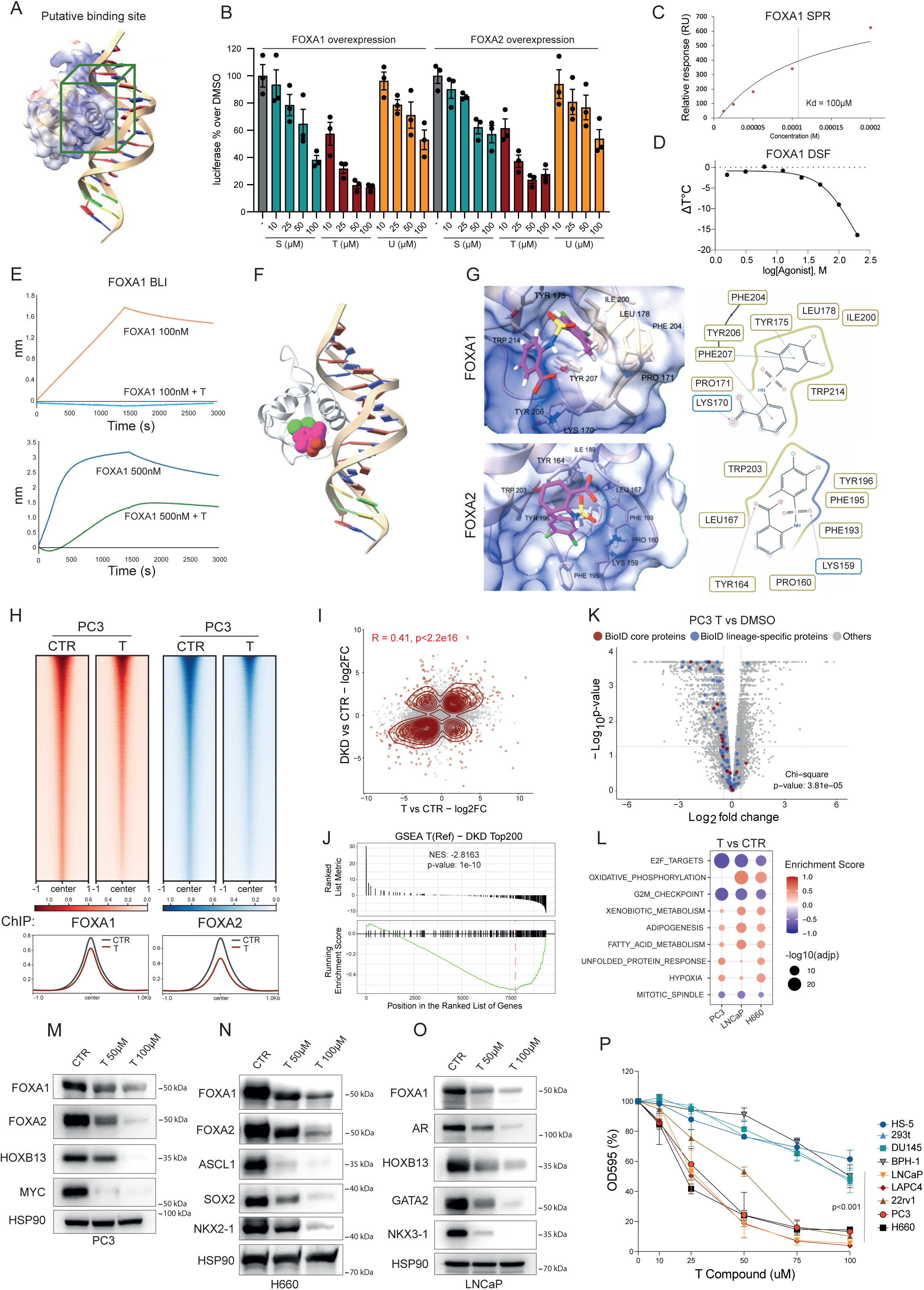
Small molecules disrupt the FOXA1/2 function. **(A)** Structure of the forkhead domain of FOXA2 with DNA highlighting the region by a green cube considered the reference binding site in the virtual screening calculations. **(B)** Luciferase validation assay shows the percent reduction of FOXA1- or FOXA-2-induced activity by the indicated concentrations of compounds S, T, and U in 293T cells over 24 hours (see Figure S6B). **(C)** Results of the Surface Plasmon Resonance (SPR) binding for the T compound using the immobilized recombinant forkhead domain of FOXA1. The dissociation constant (KD) is calculated by a steady state model and resulted to be 100µM. **(D)** Influence of different concentrations of T compound on the melting temperature of the recombinant FOXA1 forkhead domain measured by Differential Scanning Fluorimetry (DSF). **(E)** Ability of the FOXA1 forkhead domain to bind an immobilized double-strand DNA sequence in the presence and absence of 100µM of compound T by Bio-Layer Interferometry (BLI). **(F)** Graphical representation of how the T compound could impede the binding of FOXA1 to DNA. The picture was obtained by superimposing the docking solution with the FOXA1 forkhead domain. **(G)** 3D structure of the T-FOXA1 or T-FOXA2 complex (upper and lower part, respectively) inferred by unbiased docking calculations. Interaction residue is represented as a stick, while the protein structure is represented as a cartoon. The protein surface is colored by electrostatic potential. For the sake of clarity, the protein carbon atoms are depicted in grey while the T carbon atoms are in magenta. On the left are the corresponding 2D structures. **(H)** ChIPseq heatmaps and global profiles of PC3 control cells and cells treated for 1 day with 100 µM of compound T using one replicate. **(I)** Scatter plot comparing gene expression changes between PC3 cells with double knockdown of FOXA1/2 (DKD vs CTR) and cells treated with 100µM of T compound for two days (T vs CTR). Pearson correlation coefficient and corresponding p-values are indicated for genes with significant changes. **(J)** Corresponding gene set enrichment plot of top 200 downregulated genes by compound T in FOXA1/2 double-knockdown (T(ref) – DKD Top200) cells. Significance indicated by p-value. **(K)** Volcano plot of mass-spectrometry data of PC3 cells treated with 100µM compound T for 2 days. The top 100 BioID-enriched proteins are labeled, and the significance of enrichment in downregulated proteins is indicated by Chi-square analysis and corresponding p-value. **(L)** Corresponding gene set enrichment analysis of protein abundance changes in LNCaP, PC3, and H660 cells treated with T compound versus control (T vs CTR). The FDR q-values indicate the significance of the indicated gene sets. **(M-O)** Immunoblot analysis of PC3 (M), H660 (N), and LNCaP (O) cells probed for the indicated proteins after 2 days of incubation with compound T at the indicated concentrations. The corresponding analysis for FOXA1/2 knockdown in PC3, H660, and LNCaP cells is shown in Figures 3E, G, & S14A, as well as see Figure S14B-D for corresponding mRNA changes, respectively. **(P)** Dose-response curves for the indicated concentrations of T compound in the indicated cell lines (see also Figures 1B & S14E, F). One-way analysis of variance (ANOVA) test and p-value have been used to test for cell line susceptibility across cell lines with high or low expression of FOXA1 and/or FOXA2.

Subsequently, we tested whether compounds S, T, U, and the inactive compound Q bind to the recombinant forkhead domain of FOXA1 and FOXA2 by surface plasmon resonance (SPR). While compound Q did not physically interact with the FOXA1 forkhead domain, we observed increasing binding with compounds U, S, and T in line with their capacity to reduce the luciferase signals (Figures 4B & S11A). The binding of T was also ascertained for FOXA2 (Figure S11B). To confirm the SPR results using an orthogonal technique, we performed differential scanning fluorimetry (DSF). Notably, we observed a concentration-dependent reduction in the melting temperature of the FOXA1 and FOXA2 forkhead domain upon treatment with compound T (Figures 4D & S11C), indicative of a destabilizing effect on the protein’s folded structure. In line with this, the presence of compound T also disabled the FOXA1 and FOXA2 forkhead domain from binding to an immobilized double-strand DNA oligonucleotide by biolayer interferometry (BLI) (Figures 4E & S11D) ^31^.

In addition, we tested if compound T may also affect the binding between different nucleotide sequences and the recombinant forkhead domains of FOXA1 and FOXA2 using EMSA. Indeed, compound T disrupted the binding of all three nucleotide sequences (i.e. C, PAL, and 2C) to either the forkhead domains of FOXA1, FOXA2, and FOXA1/FOXA2 (Figures S11E-H). In agreement with the DSF results, the compound disabled binding regardless of whether T was added early on to the mix of DNA and forkhead domain (PRE) or later upon complex formation (POST) (Figures S11F-H).

Having secured a direct biophysical interaction and consequences on protein folding, we performed unbiased virtual docking on the entire forkhead domains of FOXA1 and FOXA2 to identify putative binding sites ^32–34^. In both instances, DynamicBind1, a newly developed AI-driven docking software capable of performing blind docking without predefined binding site requirements, identified that the aromatic moiety of T compound, featuring two chlorine atoms and a methyl group, occupied a highly conserved hydrophobic cavity within the target area selected for virtual screening (Figures 4A, G). The sequence and structural alignment showed that all residues within this cavity are conserved, except for a single substitution: tyrosine at position 206 in FOXA1 is replaced by a phenylalanine in FOXA2. The benzylic acid and sulfonamide moieties engage in electrostatic interactions with a lysine residue (Lys170 in FOXA1 and Lys159 in FOXA2) located on the protein surface. These lysine residues may play a crucial role in stabilizing the protein-DNA complex, positioned between two phosphate groups of the DNA backbone. Conceivably, the interaction of compound T with this cavity may perturb the hydrophobic core of the forkhead domain and, thereby, promote protein unfolding (Figure 4D).

We next validated if compound T may also disrupt the chromatin occupancy of FOXA1/2 in cells by ChIPseq. In LNCaP cells, compound T reduced FOXA1 chromatin binding (Figure S12A). Interestingly, motif enrichment analysis revealed that DNA binding motifs of lineage-specific transcription factors co-occupied in the BioID experiment (i.e., AR, HOXB13) were depleted while there was an increase in AP1 binding sites (bZIP), suggesting a partial rewiring of FOXA1 to other sites of the genome (Figures 3A, S12B-D & table S4_1). In PC3 cells, both FOXA1 and FOXA2 chromatin occupancy was also reduced by compound T (Figure 4H). Like LNCaP, we observed a rewiring of both FOXA1 and FOXA2 chromatin occupancy from sites co-occupied by lineage-specific transcription factors in the BioID experiment (i.e., HOXB13, AP1 (bZIP)), to ETS binding sites whose corresponding transcription factors (i.e., ETV4) were barely detected by BioID (Figures 3A, S12E-H & table S4_1). In aggregate, compound T reduced overall FOXA1/2 chromatin occupancy and rewired FOXA1/2 from DNA motifs heavily co-bound by lineage-specific oncogenic transcription factors to other areas of the chromatin.

To further interrogate the specificity of compound T, we compared the mass spectrometry data of PC3 cells treated with compound T with the corresponding data of FOXA1/2 knockdown. While the first principal component (PC) showed a major difference between the two data sets related to lentivirus-mediated transfer of the short hairpin knockdown constructs (i.e., interferon alpha and gamma signaling), the PC2 revealed coordinate changes of compound T with the single- and more so double-knockdown related to cell cycle and proliferation (Figures S13A, B & table S4_2). Moreover, mRNA changes caused by compound T correlated significantly with the double-knockdown of FOXA1/2 (Figure 4I & table S4_3). Because the knockdown data set was heavily influenced by virus-related interferon signaling, we derived a gene signature of PC3 cells treated with compound T and interrogated for enrichment in cells with FOXA1/2 double knockdown (Figure 4J). Indeed, the gene set was highly enriched in the latter, further suggesting at least some level of target specificity.

Because the joint loss-of-function of FOXA1 and FOXA2 substantially decreased the protein abundance of chromatin interactors, we wondered if the T compound would also phenocopy this behavior. Indeed, the T compound significantly reduced BioID interactors in PC3, H660, and LNCaP cells (Figures 4K & S13C-G). Consistent with the previous findings, the gene set enrichment analysis of the mass-spectrometry data revealed a decrease in signatures related to cell proliferation, while an upregulation of signatures related to unfolded protein response and xenobiotic metabolism may be directly attributed to the small molecule exposure (Figure 4L & table S4_4). Finally, we validated by immunoblotting that the reduction in protein abundance of key lineage-specific oncogenic transcription factors by compound T mimicked the ones of corresponding knockdown experiments (Figures 3E, G, 4M-O & S14A). Corresponding qPCR analysis revealed that protein expression changes were again in part paralleled by mRNA expression changes (Figures S8H, I & S14B-D).

Given these findings, we wondered if compound T selectively suppressed the clonogenic growth of cell lines expressing FOXA1 or FOXA1/2. Indeed, all prostate cancer cells exhibiting either FOXA1 or FOXA1/2 were significantly more sensitive to compound T than cells with low or no expression (Figures 4P & S14E, F). Moreover, sensitivities were largely comparable among FOXA1/2-expressing cancer cells regardless of AR expression status, implying that pharmacologic inhibition of FOXA1/2 may be an effective therapeutic strategy for targeting tumor cells at various stages of disease progression that includes AR-driven prostate cancer and various subtypes of lineage plasticity cancers.

## DISCUSSION

Our data indicate an unexpected collaboration between FOXA1 and FOXA2 in maintaining core components of the AR transcriptional machinery, even if AR has been lost in prostate cancer through lineage plasticity. While earlier work had shown that FOXA1 functions are also maintained during disease progression ^24^, we uncover that FOXA1/FOXA2 are jointly critical in enabling oncogenic lineage-specific transcription factors in various settings.

As a result, co-targeting FOXA1 and FOXA2 provides a powerful approach towards the collapse of oncogenic signaling in prostate cancer regardless of the specific subtype. We describe here that chromatin interaction proteins related to FOXA1 and FOXA2 as well as lineage-specific transcription factors are largely reduced in protein abundance upon double-knockdown of both and upon treatment with compound T. While in some cases corresponding gene expression data suggests a regulation at the transcriptional level by FOXA1/2, a disconnection between protein abundance and mRNA expression may point towards changes in protein stability. Conceivably, DNA/chromatin-interacting proteins may undergo degradation if not tethered to their functional environment ^35,36^. In agreement, degradation-related ubiquitin ligases and proteasome subunits have been specifically found in the soluble fractions of transcription factors that are not bound to chromatin ^37^. Alternatively, or in addition, post-translational modifications, such as SUMOylation ligases may also be involved as well ^38^.

Given the established function of FOXA1 in enabling estrogen receptor signaling, some of our findings may be translatable to hormone-driven breast cancer as well ^11,15^. In addition, a collaboration of FOXA1 and FOXA2 has also been described in lung adenocarcinoma ^39^, and FOXA2 is a selective dependency in small-cell lung cancer in analogy to NEPC (https://depmap.org). While a recent publication describes covalent small molecule modifiers of FOXA1 ^40^, we provide here, to the best of our knowledge, the first proof of concept that small molecule inhibitors can effectively exploit such dependencies.

While these are promising suggestions, targeting FOXA1/2 may have a potentially important on-target toxicities. Mice deleted of FOXA1 and FOXA2 reach birth but subsequently die because of metabolic derailment ^41^. Moreover, co-deletion of multiple FOXA members leads to severe liver damage ^42^. That said, only the development of more potent and selective compound derivatives and their testing in clinical trials may reveal if there is an exploitable therapeutic window for targeting FOXA1/2 in patients with cancer.

### Limitations of the study

While our key chromatin-related findings (e.g., co-localization of FOXA1/2 in AR-negative models) recapitulate across different cellular settings, a shortcoming of our ChIPseq data is the availability of one replicate in some settings (e.g., AR-negative cell lines). Moreover, the compound T is at this stage a tool compound with currently low affinity, activity, and, as a result, likely also low selectivity. For this reason, we have at this point not pursued further experiments in mouse models in vivo.

## RESOURCE AVAILABILITY

### Lead contact

Requests for further information and resources should be directed to and will be fulfilled by the lead contact

### Materials availability

This study did not generate new unique reagents.

### Data and code availability

- Data: The original bulk RNA-seq and ChIP-seq data have been deposited at GEO: GSE281794 (token: udapqwgyjlodder)
- Data: The original mass spectrometry data have been deposited at PRIDE: PXD057803 (token: YM110jNqgndZ)
- Code: The custom R and bash scripts used for data analysis and visualization in this study is available in the Github repository and accessible at https://github.com/211JptLab/Formaggio_FOXA2 with a unique identifier https://doi.org/10.5281/zenodo.16744575.
- All other items: Additional resources are referenced in the Key Resources Table.
- Any additional information required to reanalyze the data reported in this paper is available from the lead contact upon request.

## ACKNOWLEDGMENTS

We thank all laboratory members for their scientific discussions. The pGL-6xFBS-Luc FOXA1 reporter plasmid has been a kind gift of Charles Sawyers and Wouter Richard Karthaus. The following funding sources to J.P.T. have supported this work: Swiss National Science Foundation (310030_197810, CRSII5_186405), Krebsliga Schweiz (KFS-5136-08-2020), Fondazione Dr. Ettore Balli, Swiss Cancer Foundation, Fondazione Gustav & Ruth Jacob, Lega Ticinese Contro il Cancro, and the Foundation Nelia and Amadeo Barletta.

## AUTHOR CONTRIBUTIONS

N.F. and J.P.T. originally developed the concept, further elaborated, and designed the experiments. Y. U., D.B., and F.C. contributed to the experiments. A.C. and J.S. performed the virtual drug screen on the forkhead binding domain of FOXA2, identified hit compounds, and performed SPR, DSF, BLI, and virtual docking analyses. M.P. and R.G. performed MS experiments related to BioID, and G.T. and N.F. analyzed the data. R.S. and D.C.C. provided reagents for Bio-ID and advice. G.C. and M.B. generated and analyzed the single-cell RNA sequencing atlas. A.R. generated RNA-seq and ChIP-seq data, and C.L. and G.T. subsequently analyzed the data. J.P.T provided funding and resources. J.P.T., N.F., G.T., J.S., and A.C. interpreted the data and wrote the paper.

## DECLARATION OF GENERATIVE AI AND AI-ASSISTED TECHNOLOGIES

No AI-assisted technologies have been used for the generation of the manuscript apart from AI-assisted docking of T with the recombinant forkhead domains of FOXA1 and FOXA2 (Figure 4G).

## SUPPLEMENTAL INFORMATION

Document S1. Figures S1–S14 and table S1

Supplementary tables from table S1.1 to table S4.4

## STAR METHODS

### KEY RESOURCES TABLE

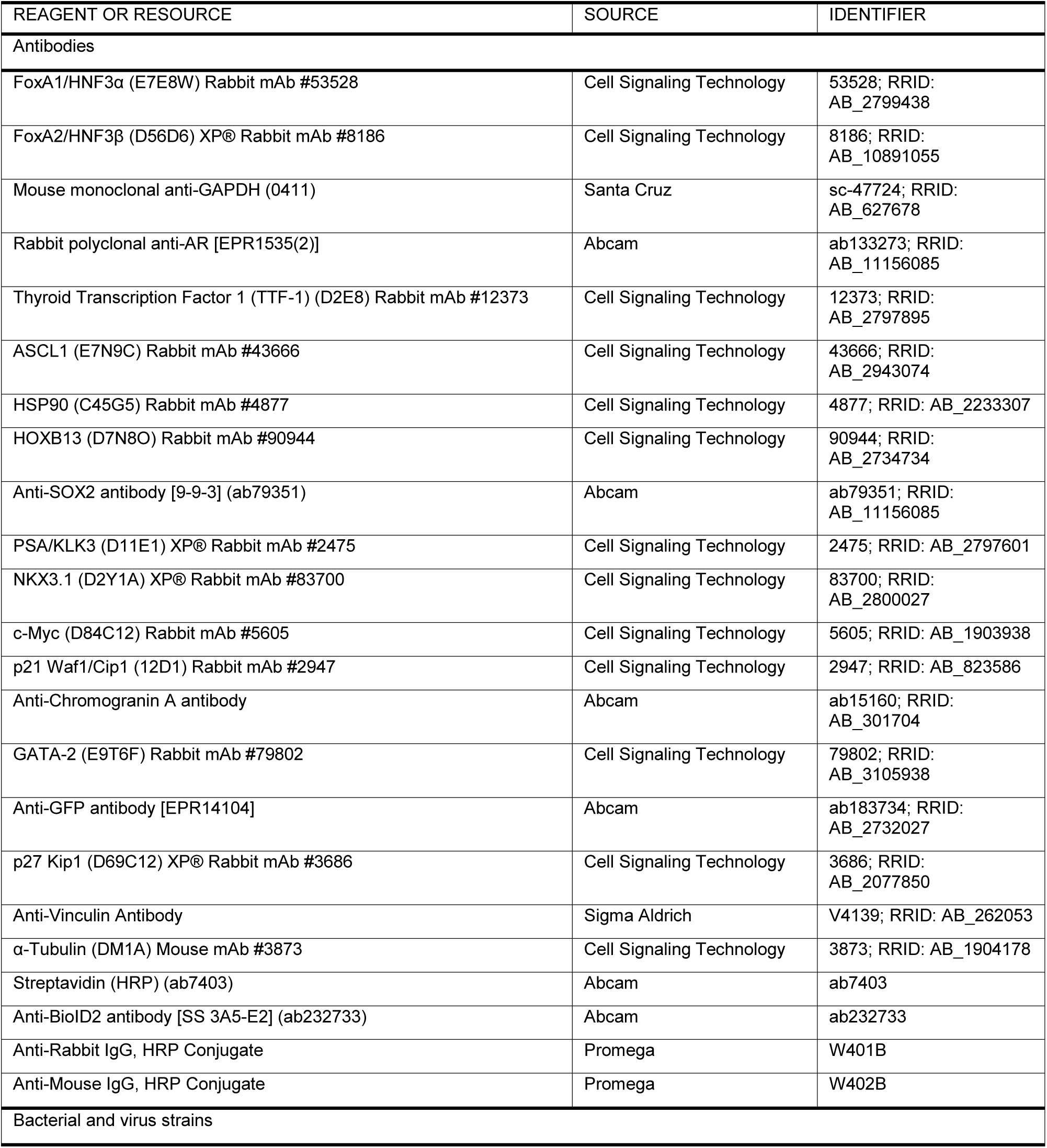
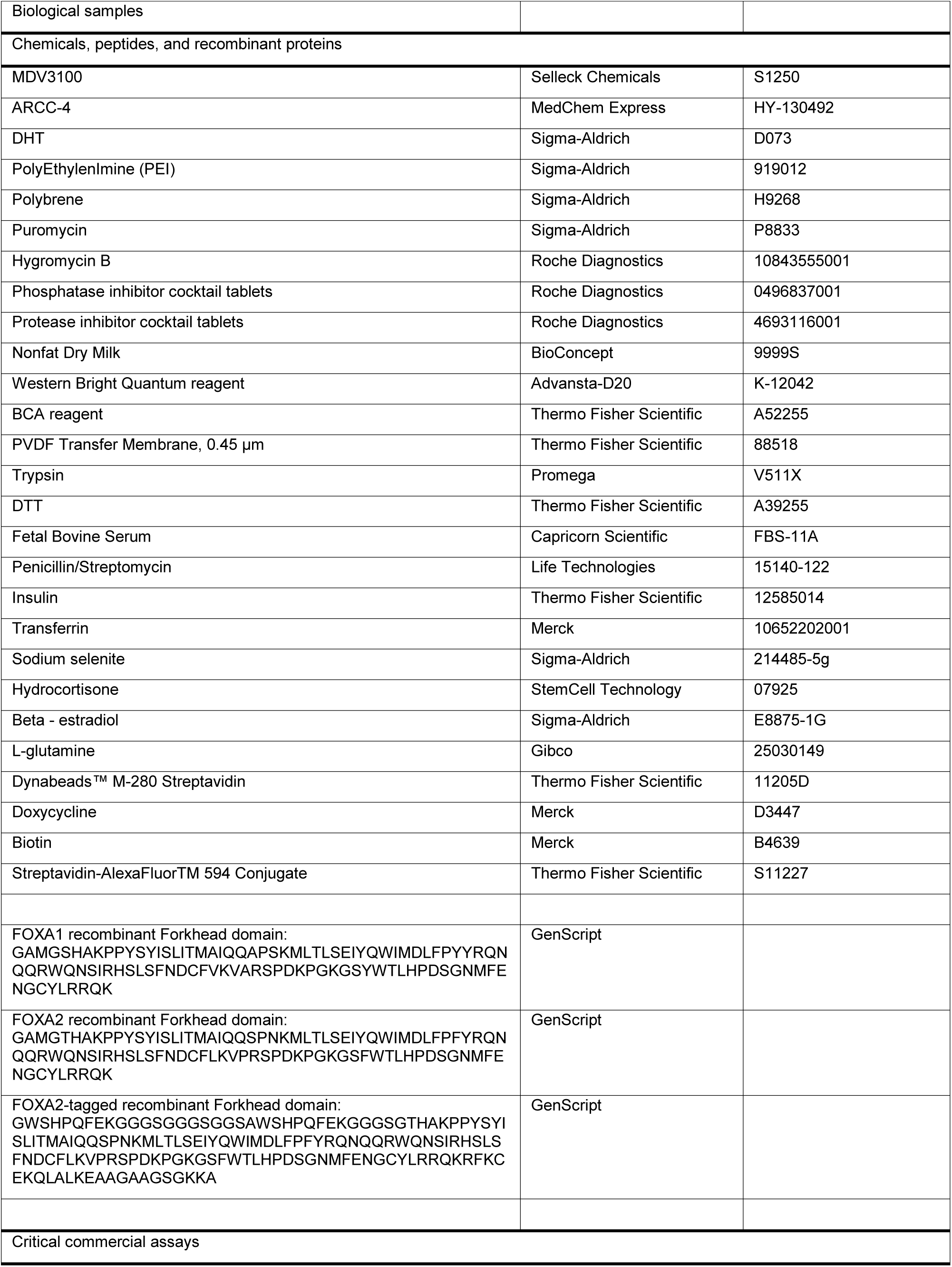
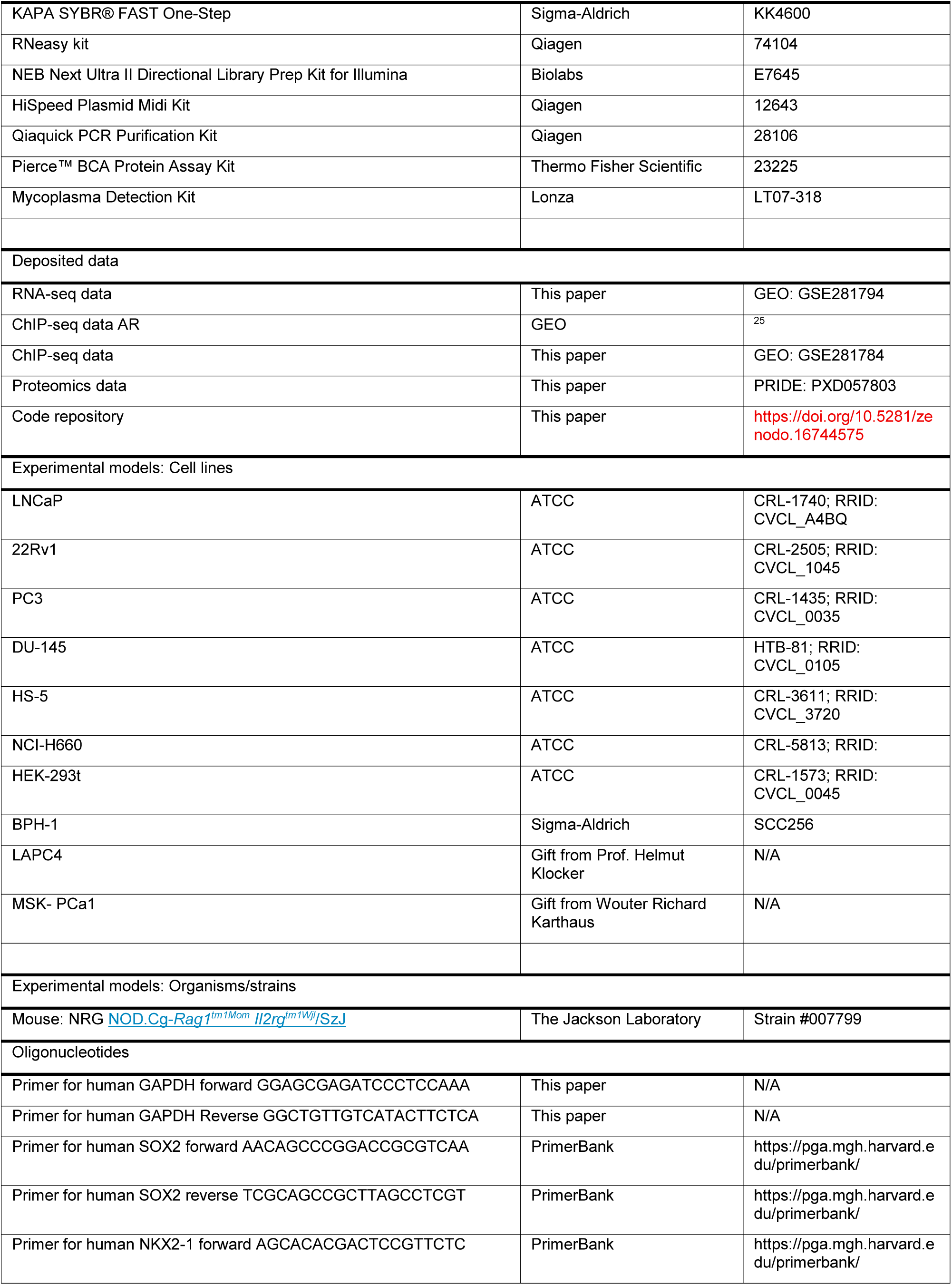
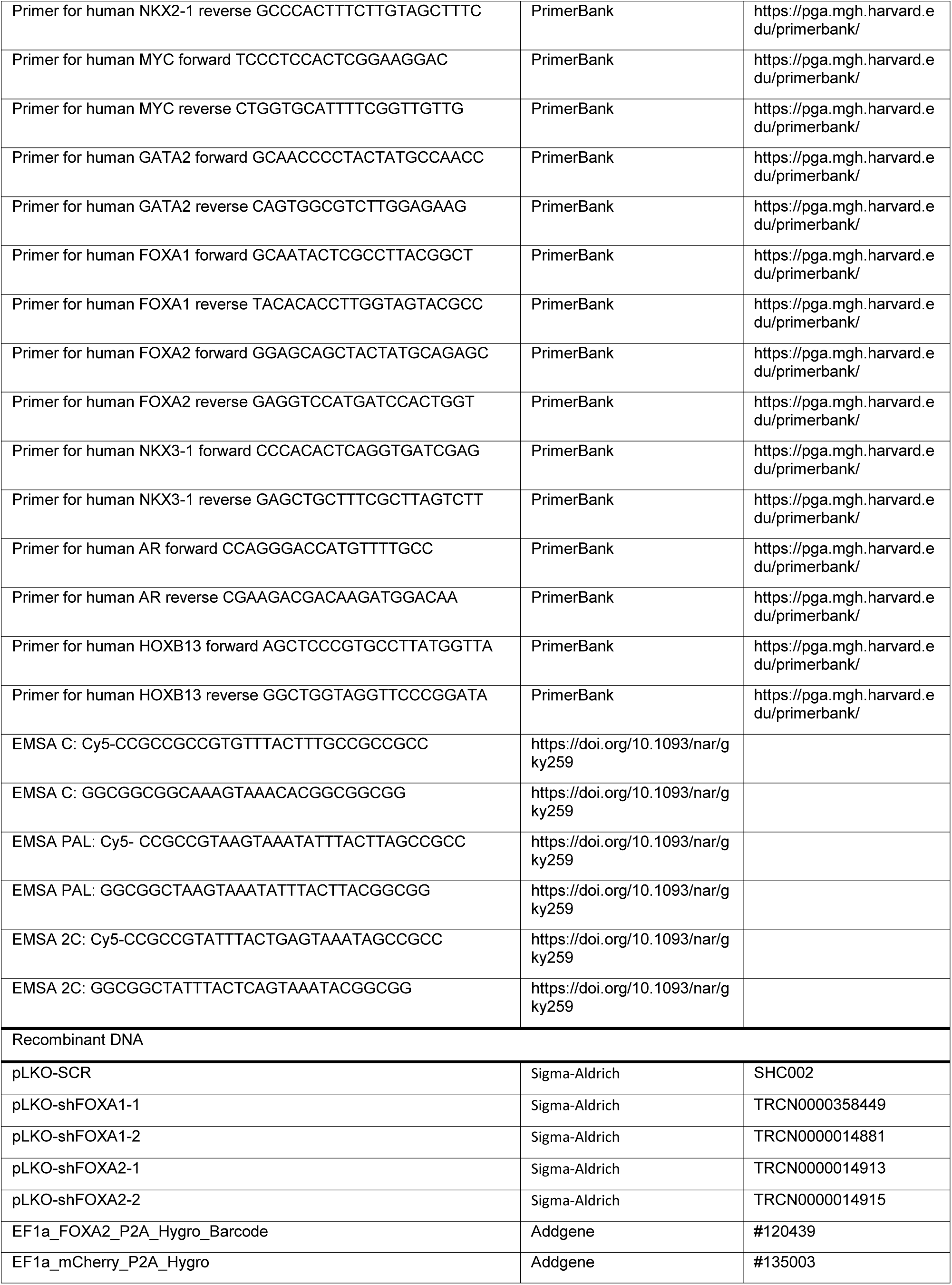
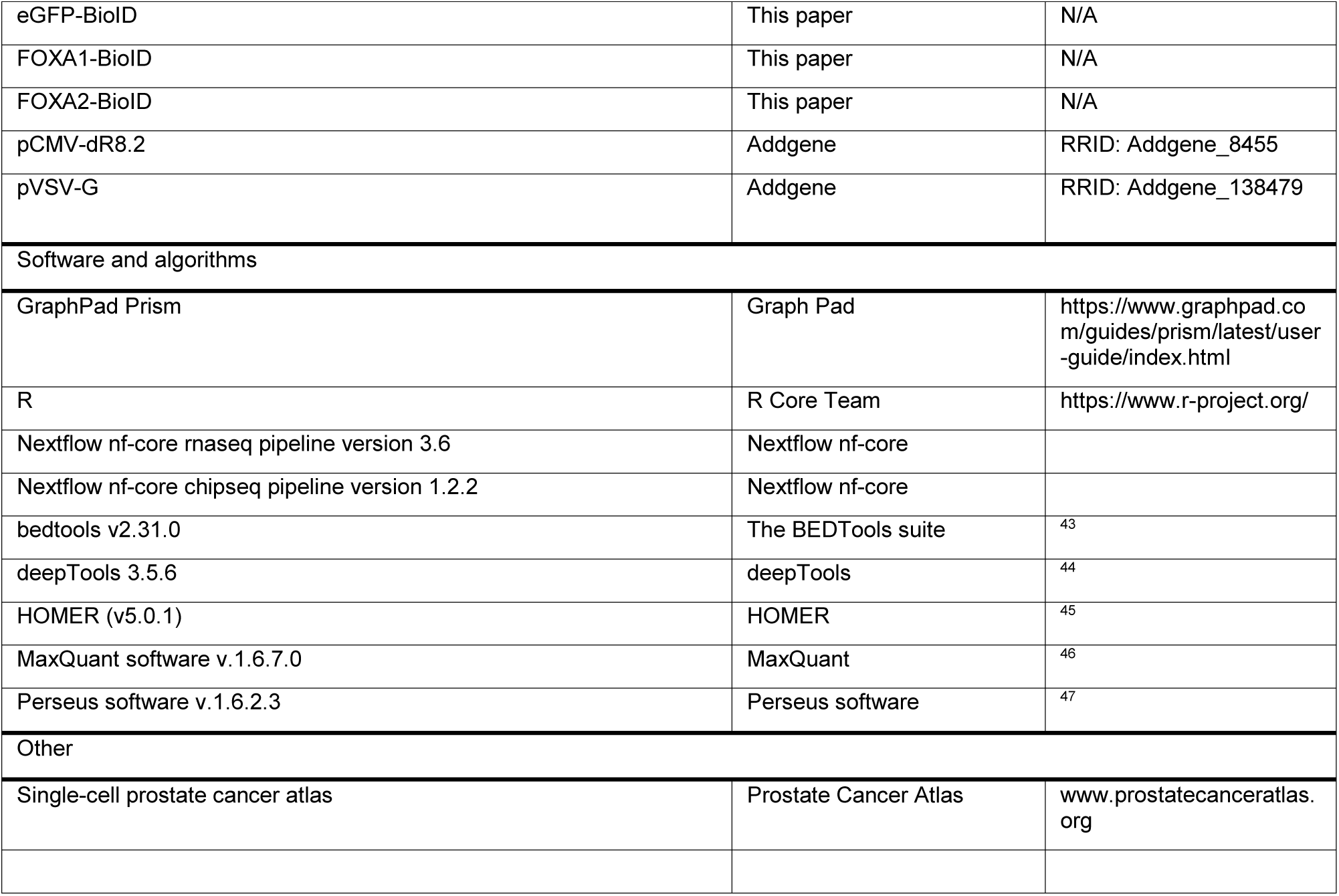

### EXPERIMENTAL MODEL AND STUDY PARTICIPANT DETAILS

#### Cell, PDX and organoid lines

LNCaP, 22rv1, PC3, DU145, HS-5, HEK 293T and H660 cell lines were purchased from ATCC (American Tissue Culture Collection) (Manassas, USA). The BPH-1 cell line was purchased from Sigma-Aldrich. The LAPC-4 cell line was a gift from Prof. Helmut Klocker. The supernatant of all cell lines was routinely tested (once per month) using the MycoAlertTM Mycoplasma Detection Kit (Catalog #: LT07-318 Lonza). All cell lines tested negative for Mycoplasma infection. LuCaP 145.2 PDX was provided by Dr. Eva Corey. MSK-PCa1 human organoid line was provided by Wouter Richard Karthaus.

#### Animal experiments

LuCaP 145.2 mouse model was established from specimens acquired at either radical prostatectomy or autopsy, implanted and maintained by serial passage by implanted Matrigel-embedded tissue tumor fragments into NRG male mice. MSK-PCA1 PDOX model was established from specimens acquired at either radical prostatectomy or autopsy, implanted and maintained by serial passage by implanted Matrigel-embedded tissue tumor fragments into NRG male mice. All animal experiments were conducted following a protocol that was approved by the Swiss Veterinary Authority/Board (TI-42-2018, TI-30-2021, TI-70-2024,) and received approval from the ethical committee of the Institute of Oncology Research (IOR). For all in vivo studies, we used male NRG (NOD-Rag1null IL2rgnull, NOD rag gamma) mice between 6-8 weeks old.

### METHOD DETAIL

#### Cell culture

The LNCaP, 22rv1, PC3, DU145 cell lines were cultured in RPMI 1640 medium (Cat 21875-24, Gibco) supplemented with 10% Fetal Bovine Serum (FBS-11A Capricorn Scientific) and 1% Penicillin/Streptomycin (15140-122 Life Technologies) with 5% CO2 at 37°C. The LAPC-4 cell line was cultured in RPMI 1640 medium (Cat 21875-24, Gibco) supplemented with 10% Fetal Bovine Serum (FBS-11A Capricorn Scientific), 1nM of DHT and 1% Penicillin/Streptomycin (15140-122 Life Technologies) with 5% CO2 at 37°C. The HEK 293T and HS-5 cell lines were cultured in DMEM medium (Cat 2688-133, Gibco) supplemented with 10% FBS and 1% µg/ml Penicillin/Streptomycin with 5% CO2 at 37°C. H660 cells medium was generated accordingly to ATCC protocol in RPMI 1640 medium (Cat 21875-24, Gibco) 0.005 mg/ml Insulin (Thermo Fisher Scientific 12585014), 0.01 mg/ml Transferrin (Merck 10652202001), 30nM Sodium selenite (Sigma 214485-5g), 10 nM Hydrocortisone (StemCell Technology 07925), 10 nM beta-estradiol (Sigma E8875-1G), extra 2mM L-glutamine (Gibco 25030149), 5% fetal bovine serum (FBS-11A Capricorn Scientific), 1% µg/ml Penicillin/Streptomycin with 5% CO2 at 37°C. BPH-1 cells were cultured in RPMI1640 medium (Cat 21875-24, Gibco) with 20% FBS (FBS-11A Capricorn Scientific), 30nM Sodium selenite (Sigma 214485-5g), 0.005 mg/ml Insulin (Thermo Fisher Scientific 12585014), 1nM DHT and 1% Penicillin/Streptomycin (15140-122 Life Technologies) with 5% CO2 at 37°C.

#### PEI-mediated transfection and lentiviral infection

The HEK 293T cells were transfected with a vector for each shRNA / overexpression plasmid, packaging, and envelope vector using the PolyEthylenImine (PEI) method. The HEK 293T cells were seeded in a 100 mm culture dish (4×106 cells/plate) and incubated overnight at 37°C in a 5% CO2 humidified atmosphere. After 24 hours, the vector plasmid (3 µg), packaging plasmid (pCMV-dR8.2; 2.7 µg), and envelope plasmid (pVSV-G; 0.7 µg) were mixed in Opti-MEM™ I Reduced Serum Medium (300 µL/10 cm culture dish; Cat 31985-070, Gibco) and with 1.25 mM PEI (Sigma-Aldrich 919012) solution (ratio µL PEI: µg DNA 4:1). The DNA/PEI mixture were incubated for 15 minutes at room temperature and added to the HEK 293T cell supernatant. After 48 hours of transfection, the viral supernatants were collected and filtered through a 0.45µm filter. The different cell line is then incubated with the viral supernatant and 8µg/ml Polybrene (H9268 Sigma) for 72 hours and then selected with 2 µg/ml puromycin (P8833 Sigma) or 100 µg/ml Hygromicyn B (10843555001, Roche Diagnostics) for one week.

#### Plasmids

The pLKO-SCR (SHC002Sigma-Aldrich), the pLKO-shFOXA1-1 (TRCN0000358449, Sigma-Aldrich), the pLKO-shFOXA1-2 (TRCN0000014881, Sigma-Aldrich), the pLKO-shFOXA2-1 (TRCN0000014913, Sigma-Aldrich) and the pLKO-shFOXA2-2 (TRCN0000014915, Sigma-Aldrich) were purchased from Sigma. The EF1a_FOXA2_P2A_Hygro_Barcode (Addgene #120439) and the EF1a_mCherry_P2A_Hygro (Addgene #135003) were purchased by Addgene. Dr. Raffaella Santoro provided BioID-nuclear-eGFP plasmid. FOXA1-BioID and FOXA2-BioID plasmids were produced in house cloning FOXA1 and FOXA2 human cDNA from Genewiz (Azenta Life Science) instead of eGFP.

#### Antibodies for immunoblot and immunoprecipitation

The primary antibodies used are anti-FOXA1 (E7E8W, CST), anti-FOXA2 (D56D6, CST), anti-GAPDH (0411-Santa Cruz), anti-AR (133273 Abcam), anti-NKX2-1 (D2E8, CST), anti-ASCL1 (E7N9C, CST), anti-HSP90 (C45G5, CST), anti-HOXB13 (D7N80, CST), anti-SOX2 (ab79351, Abcam), anti-PSA (D11E1, CST), anti-NKX3-1 (D2Y1A, CST), anti-MYC (D84C12, CST), anti-p21(12D, CST), anti-CHGA (ab15160, Abcam), anti-GATA2 (E9T6F, CST), anti-GFP (EPR14104, Abcam), anti-p27(D69C12, CST), anti-VCL (SAB14004522, SIGMA), anti-Alpha Tubuline (DM1A, CST), streptavidin-HRP (ab7403, Abcam), anti-BioID2 (ab232733, Abcam). Snap-frozen cellular pellet was lysed using RIPA Buffer supplemented with a cocktail of phosphatase inhibitors (4906845001 Roche) and protease inhibitors (5892953001 Roche). The protein concentration was determined using a BCA reagent (A52255 Thermo Fisher Scientific). 30-50 µg of whole protein lysate was separated on 8-12% SDS-polyacrylamide gels and transferred onto PVDF membrane (88518 Thermo Fisher Scientific). The membranes were first blocked with 5% milk in Tris Buffered Saline with Tween 20 (TBST) for 30 minutes at room temperature. After that, they were incubated with primary antibodies overnight at 4°C. Then, they were incubated with secondary antibodies (anti-rabbit IgG HRP W401B and anti-mouse IgG HRP W402B Promega) for 1 hour at room temperature. The protein bands were visualized using the western bright quantum reagent (K-12042-D20 Advansta) and quantified using the Fusion Solo IV LBR system.

#### Proliferation assay

PC3 (1k) and LNCaP (4k) were seeded in 96-well plates, and the day after, they were treated or not with Enzalutamide 10µM and put inside Incucyte SX3 (Sartorius) for 7 days (one scan every 6 hours). Confluence was evaluated in fold change over time 0 and normalized over the specific control of the experiment. For H660 cells, 3D methylcellulose assay was performed with Incucyte SX3 (Sartorius) using spheroid module. 40k cells were seeded in a single cell suspension in full H660 medium with 40% of MethoCultTM (StemCell Technologies) and scans were taken every 6 hours for 15 days. The dimension of the 3D clusters was evaluated with fold change in confluence mask over time 0 and normalized over the specific control of the experiment. FOXA1 and FOXA2 double-knockdown experiments in PC3 and H660 cells were conducted as discussed previously, but the endpoints were prolonged to show the differences with the single knockdown.

#### ARCC4 resistance assay

50k LNCaP cells were seeded in each well of a 6 well plate. Once attached (the day after), cells were treated with DMSO or ARCC4 1µM. The medium was renewed with treatment twice a week. The endpoint of the experiment was after 45 days of treatment.

#### RNA extraction for RNA-seq analysis

According to the manufacturer’s guidelines, the RNA extraction was performed from the prostate cancer cell lines and fresh tumours using QIAGEN RNAeasy Mini Kit. The RNAs were processed using the NEBNext Ultraexpress RNA Library Prep Kit for Illumina (E3330S - New England BioLabs Inc.) with NEBNext Multiplex Oligos (New England BioLabs Inc.) and NEBNext® Poly(A) mRNA Magnetic Isolation Module for cDNA synthesis (E7490 - New England BioLabs Inc.) with the addition of barcode sequences and sequenced on Illumina NextSeq 2000 with single- end, 120-base-pair long reads.

#### RT-qPCR

RNA was extracted from a cellular pellet of LNCaP cells using RNeasy kit (74106 Qiagen), following the manufacturer’s guidelines. The extraction was performed under the indicated culture conditions. RT-qPCR analysis was carried out using KAPA SYBR® FAST One-Step (KK4600 Sigma) following the manufacturer’s protocol. The primer sequences were obtained from PrimerBank (http://pga.mgh.harvard.edu/primerbank/index.html). The housekeeper gene used was GAPDH. The qPCR analysis was performed using the 2-ΔΔ Ct method.

#### BioID experiments

For BioID pulldown assay, 50 × 106 LNCaP, PC3 or H660 cells were plated. The day after, different concentration of Doxycycline (from 100ng/mL to 1µg/mL, D3447, Merck) were used to induce at the same levels different BioID constructs (GFP-BioID, FOXA1-BioID and FOXA2-BioID). Together with Doxycycline, 50µM of Biotin (B4639, Merck) was added to the medium. 24 hours later cells were collected. After two PBS washes, nuclear separation was performed using 20mM Tris-HCl, 10 mM NaCl, 3 mM MgCl2. Cytosol was discarded and nuclei were washed with ice cold PBS two times. Only for H660 cells the whole cell lysate was used for streptavidin pulldown since nuclear separation was sufficiently accurate and reproducible. Nuclei were lysed in complete RIPA buffer and sonicated using Bioruptor Plus (BIOSENSE) (high power, 30 cycles, 30 sec ON 30 sec OFF). After BCA protein quantification, 4mg of nuclear protein were loaded in 4mL volume of RIPA buffer and incubated rotating overnight with 50µL of Dynabeads™ M-280 Streptavidin (11205D, ThermoFisher) magnetic beads. After the pulldown, several washes were performed: 2 in RIPA buffer, 2 in SDS 2% buffer, 4 in Litium High Salt Buffer (Tris pH 8.0 50mM, LiCl 500mM, EDTA 2mM), 4 in Na High Salt Buffer (Tris pH 8.0 50mM, NaCl 500mM, EDTA 2mM), 2 in Low Salt Buffer (Tris pH 8.0 50mM, 150mM NaCl). Magnetic beads were made dry and snap frozen in liquid nitrogen. For western blot analysis, beads were eluted with 4x Laemmli Buffer 95°C.

#### Liquid chromatography-tandem mass spectrometry (LC-MS/MS)

Protein extraction and enzymatic digestion: For the proteomes preparation, a cell pellet of approximately 1 × 106 cells was washed twice in phosphate-buffered saline (PBS), flash frozen and stored at −80°C. Cell lysis and protein extraction were performed in 8M urea in 50 mM ammonium bicarbonate (ABC) by sonication (Bioruptor, Diagenode, 15 cycles, 30s on, 30s off, high mode). Proteins were reduced with 10 mM dithiothreitol for 20 minutes at room temperature and alkylated with 50 mM iodoacetamide for 30 minutes at room temperature. Digestion was carried out in 8M urea in 50 mM ABC for 2 hours at room temperature with LysC (Wako Fujifilm, 1:100 w/w). After diluting the digestion buffer to 2M urea with 50 mM ABC, trypsin (Promega, 1:100 w/w) was added for overnight digestion at room temperature. Digestion was halted by adding acetonitrile (ACN) to 2% and trifluoroacetic acid (TFA) to 0.3% and the samples were cleared by centrifugation for 5 minutes at maximum speed. The digested peptides were purified with C18 StageTips ^48^, and eluted with 80% ACN, 0.5% acetic acid. Finally, the elution buffer was dried by vacuum centrifugation and purified peptides were resuspended in 2% ACN, 0.5% acetic acid, 0.1% TFA for LC-MS/MS analysis. The BioID samples were prepared with the same procedure with minor modifications. In brief, beads were resuspended in 8M urea in 50 mM ABC, and proteins were reduced with 10 mM dithiothreitol for 60 minutes at 37°C shaking and alkylated with 50 mM iodoacetamide for 30 minutes at room temperature shaking. On-bead sequential digestion with 1 µg of LysC and trypsin was carried out at 37°C shaking for 2 hours and overnight, respectively, and peptides desalting and purification were performed.

#### LC-MS/MS analysis

Proteomes were analyzed on a timsTOF HT mass spectrometer (Bruker) coupled online via a nanoelectrospray source (Captive spray source, Bruker) to a nanoElute2 HPLC system (Bruker). 500 ng of peptides per sample were loaded in water/0.1% formic acid into a 75 µm inner diameter, 25 cm long column in-house packed with ReproSil-Pur C18-AQ 1.9 µm resin (Dr. Maisch HPLC GmbH) kept at 50°C in a column oven, and eluted over a 60-min linear gradient between 2 and 35% ACN/0.1% formic acid at a flow rate of 300 nl/min. The mass spectrometer was operated in a data-independent (DIA)-PASEF mode with an accumulation and ramp time of 100 ms, covering with 21 mass steps, 25 Da wide, and 1 mobility window, a mass range from 475 to 1000 Da and a mobility range from 0.85 to 1.27 Vs/cm2, with an estimated cycle time of 0.95 s. Collision energy was ramped linearly from 20 eV at 0.60 Vs/cm2 to 59 eV at 1.60 Vs/cm2.

BioID experiments were analyzed on a Q Exactive HF mass spectrometer (Thermo Fisher Scientific) coupled online via a nanoelectrospray source (Thermo Fisher Scientific) to an EASY-nLC 1200 HPLC system (Thermo Fisher Scientific). 1 µg of peptides per sample was loaded in buffer A (0.1% formic acid) into a 75 µm inner diameter, 50 cm long, packed in-house column, kept at 50°C, and eluted over a 150-min linear gradient of 5 to 30% buffer B (80% ACN, 0.1% formic acid) at a flow rate of 250 nl/min. The instrument was operated in a data-dependent (DDA) mode with a survey scan range of 300-1,650 m/z, resolution of 60,000 at 200 m/z, maximum injection time of 20 ms, and AGC target of 3e6. The top 10 most abundant ions with charges 2 to 5 were isolated with a 1.8 m/z isolation window and fragmented by higher-energy collisional dissociation (HCD) at a normalized collision energy of 27. MS/MS spectra were acquired with a resolution of 15,000 at 200 m/z, maximum injection time of 55 ms, and AGC target of 1e5. Dynamic exclusion was set to 30 s to avoid repeated sequencing.

#### Single-cell prostate cancer atlas (www.prostatecanceratlas.org)

For the harmonized single-cell prostate cancer atlas, the raw data from the following studies were retrieved: GSE117403, GSE133094, GSE135520, GSE137829, GSE143791, GSE153892, GSE176031, GSE181294, GSE185344, GSE193337, GSE157703, GSE210358, HDAC000500, GSE179312, GSE145838, GSE145843, and PRJNA699369. Raw FASTQ files were obtained from the original sources, aligned to the GRCh38 genome using STAR-solo (v2.7.10b), and converted to feature-barcode matrices. Each sample was imported into R (v4.0.2) as a Seurat object (v4.0.3) ^49–51^ and underwent initial quality filtering using scuttle (v1.8.1) ^52^, removing low-quality cells based on cell-level metrics (e.g., UMI counts, mitochondrial content). Doublets were estimated and removed via DoubletFinder (v2.0.3) ^53^. We applied global scaling normalization and log transformation, followed by selecting highly variable features. Subsequently, we integrated and clustered individual cells into cohesive groups. To harmonize the scRNA-seq data, we utilized deep generative models from scvi-tools ^54^, specifically variational autoencoders (v0.14.5, n_latent=30, n_layers=2, gene_likelihood=“nb”). This approach allowed us to correct for batch effects related to donor identity, sorting strategy, and library generation kits, as well as to control for unwanted sources of variation, including total counts, number of features, percent mitochondrial genes, and percent ribosomal genes (specified in ‘continuous_covariate_keys’). We used the 20 latent dimensions to determine a KNN graph for UMAP generation and leiden clustering. Granularity was assessed with clustree (v0.5.0) ^55^, selecting a clustering resolution of 1. Marker identification was conducted via MAST ^56^, applying Bonferroni correction, and only genes meeting stringent expression and significance thresholds (p_adj < 0.05, log2FC > 1) were deemed markers. Following annotation, we subsetted epithelial and malignant clusters, retaining cells from healthy, primary, and castration-resistant tumor samples. Specific biological pathway associations, including AR, neuroendocrine (NE), stem-cell-like (SCL), and WNT-signaling, were assessed using AddModuleScore ^9^. The Seurat object was converted to an anndata object, and Scanpy (version 1.9.5) toolkit was employed for visualization purposes ^57^.

#### Confocal microscopy

40k PC3 cells were plated on a 13mm coverslip glass in a 24-well plate. The day after were treated with 1µg/mL of Doxycycline (D3447, Merck) and 50µM of Biotin (B4639, Merck). 24 hours later cells were fixed with 10% formalin for 10min, washed two times with PBS and permeabilized by 0.4% Triton X-100/PBS for 15 min. Cells were blocked with 1% BSA in 1X PBS-Tween 20 (0.1%) for 60 min and then Streptavidin-AlexaFluorTM 594 Conjugate (S11227, ThermoFisher) was added 1:500 for 60 min. After PBS washes, coverslips were mounted using Fluoromount-G™ Mounting Medium, with DAPI. Images were taken with Leica TCS SP5.

#### Colony assay

30k PC3, LNCaP, 22rv1, DU145, LAPC-4, HEK 293T, HS-5 and BPH-1 were seeded in each well of a 6-well plate. The day after, cells were treated with 10µM, 25µM, 50µM, 75µM, 100µM of compound T. Endpoint was obtained when DMSO-treated cells reached a sufficient level of confluence (in a range of 2-4 weeks after the seeding). Before Crystal violet staining, full images of each well were obtained with Incucyte SX3 (Sartorius). After washing and drying of crystal violet, images were taken using Fusion Solo IV LBR system. Finally, crystal violet solution was dissolved using 33% acetic acid and absorbance (595nM) was measured with Cytation 5 Cell Imaging Multi-Mode Reader (BioTek). For H660, cells were plated in full medium (as described before) with 40% of MethoCultTM (StemCell Technologies) and the readout was taken with Incucyte SX3 (Sartorius).

#### Luciferase reporter assays

293T cells were transiently transfected using JetPrime (Polyplus) with the pGL-6xFBS-Luc (Adams et al., 2019) along with a pRL-CMV-Renilla (Renilla luciferase) internal control and the vector of interest: either pcw107 empty backbone (addgene #62511), pcw107-FOXA1 (in house cloned) or pcw107-FOXA2 (in house cloned) for each experiment. To optimize the assay the reporters were dose-response tested using different levels of pcw107-FOXA1 and pcw107-FOXA2. Compounds treatment was performed 2 hours after the transfection. Luminescence was measured 24h post-transfection using Dual-Glo Luciferase Assay System (Promega E2920) and response ratios were normalized to activity of pcw107-FOXA1 or pcw107-FOXA2 subtracting negative control activity of empty vector pcw107 (without exogenous FOXA1 or FOXA2). All results are means and standard deviations from experiments performed in biological triplicates and Firefly luciferase activity of individual wells was normalized against Renilla luciferase activity.

#### Chromatin immunoprecipitation (ChIPseq)

50 million cells were fixed with 1% formaldehyde at room temperature for 10 min and quenched. Nuclear separation was performed using 50 mM Hepes-KOH pH 7.5, 140 mM NaCl, 1 mM EDTA pH 8.0, 10% glycerol, 0.25% Triton X-100 and 0.5% NP40 and protease inhibitor (05892953001, Roche). Nuclei were washed and after lysed in lysis buffer (10 mM Tris HCl pH 8.0, 100 mM NaCl, 1 mM EDTA pH 8.0, 0.5 mM EGTA pH 8.0, 0.1% Na-deoxycholate (freshly made), 0,5% N-lauroylsarcosine Salt and protease inhibitor (05892953001, Roche). Chromatin was sonicated to 200-600 bp using a Bioruptor Plus (BIOSENSE) (high power, 30 cycles, 30 sec ON 30 sec OFF). Antibodies (5µg of FOXA1 ab23738 Abcam; 5µg of H3K27ac C15410196 Diagenode; 10 µg of FOXA2 D56D6 CST) were incubated with sonicated chromatins overnight. Subsequently 50 µl of Dynabeads protein G (Invitrogen) were added for 5 to 6 hours. Chromatins were washed with NaCl wash buffer (50 mM Hepes pH 7.8, 500 mM NaCl, 1mM EDTA, 1% Triton X 100, 0.1% Na-deoxycholate and protease inhibitor) two times for 10 min sequentially. Chromatins were washed with LiCl wash buffer (20 mM Tris-HCl pH8, 250 mM LiCl, 1mM EDTA, 0.5% NP-40 and protease inhibitor) two times for 10 min sequentially. DNA extraction was performed using QIAquick PCR Purification Kit (Qiagen).

For MSK-PCa1 and LuCaP 145.2 freshly removed PDX tumors were dissociated using gentleMACSTM tissue dissociator having small cluster of cells. After the dissociation, we filtered the cell suspension through a 100 µM cell strainer (Roche) to eliminate macroscopic tissue pieces and then centrifuged it. The cell pellet was resuspended in RPMI complete medium, the following protocol has been conducted in the very same way as for cell lines.

#### Differential Scanning Fluorimetry (DSF)

Thermal stability of the protein samples was assessed using differential scanning fluorimetry (DSF) using the Protein Thermal Shit Dye Kit. The DSF experiments were performed in a total volume of 20 µL in 96-well PCR plates. Each well contained a final concentration of 2 µg of FoxA1 forkhead domain. The reaction mixture was subjected to a thermal ramp from 25°C to 95°C at a rate of 0.05°C/sec using a qPCR machine. The experiments were performed using increasing concentration of T (from 3.12 to 200 µM). All the values are the average of the results obtained in four different wells. The Boltzmann-derived melting temperature Tm was plotted as function of the drug concentration. As a negative control, a single experiment considering a 100 mM concentration of compound Q was run. In this case, we did not observe any perturbation of Tm.

#### Surface Plasmon Resonance (SPR)

The binding interactions between T, U, S, and Q with the FoxA1 forkhead domain were studied using surface plasmon resonance (SPR). The protein was immobilized on a CM5 chip via amine coupling in acetate buffer at pH 4.5. A single-cycle kinetic method was used to inject six increasing concentrations of small molecule Q, ranging from 6.25 to 200 µM. Phosphate-buffered saline (PBS) containing 2% DMSO was employed as the running buffer to enhance solubility. Curve fitting was subsequently performed using Biacore Insight Evaluation Software (version 5.0.18).

#### Biolayer Interferometry (BLI)

A biotinylated double-stranded oligonucleotide (sequence /5BiotinTEG/GT GTC TCC TGC TCT GTC AGC AGG GCA CTG TAC TTG CTG AT and ATC AGC AAG TAC AGT GCC CTG CTG ACA GAG CAG GAG ACA C, derived from a recently released cryo-EM structure) was dissolved in Sartorius Kinetics Buffer at a concentration of 2 µg/mL and immobilized onto Streptavidin (SA) Biosensors. The binding interaction between the oligonucleotide and recombinant FOXA1 forkhead domain was evaluated at protein concentrations of 100 nM and 500 nM. The assay was conducted both with the protein alone and in the presence of compound T at a concentration of 100 µM. Non-specific signals were corrected by subtracting the signal from a sensor without immobilized biotin. All experiments were performed on an Octet® R8 system, and the resulting data were plotted and fitted using a 2:1 kinetic model in Octet® Analysis Studio (v.12.2.2.26)

#### Electrophoretic Mobility Shift Assay (EMSA)

5’ Cy5-labeled forward strand DNA oligo and the corresponding reverse complementary unlabeled strand (HPLC-purified, purchased from Integrated DNA Technologies) were first annealed in 1X annealing buffer (20 mM Tris–HCl, 50 mM MgCl2, 50 mM KCl) by heating to 95 °C for 5 min and gradual cooling at 1 °C /min to 16 °C (10μM). 100 nM of dsDNA was incubated with increasing concentrations of the FOXA1 forkhead domain protein in 1X binding buffer (20 mM Tris-HCl, pH 8.0; 0.1 mg/ml bovine serum albumin; 50 μM ZnCl2; 100 mM KCl; 10% (v/v) glycerol; 0.1% Igepal CA-630 and 2 mM β-mercaptoethanol) for 1h at 4°C and 10 µL of sample was loaded onto pre-run 12% native polyacrylamide mini-gels in 1X TAE buffer and electrophoresed for 30 min at 200 V on ice. The images were detected Typhoon biomolecular imager. Sequences of all tested DNA oligos are listed in the Key Resource Table. For heterodimer formation, a FOXA2-tagged recombinant protein with an additional 6kDa was used. When used together for heterodimerization, both proteins (FOXA1 and tagged-FOXA2) were used at 200nM. The aminoacidic sequence of all the recombinant proteins used is listed in the section Key Resource Table.

#### Virtual screening

Virtual screening calculations were conducted using the Schrodinger molecular modeling suite. Compound libraries were prepared with the LigPrep module, ensuring correct structures and protonation states. The virtual screening (VS) process was carried out with Glide employing progressively more sophisticated scoring functions: High Throughput Screening (HTS), Standard Precision (SP), and Extra Precision (XP). Finally, the top 20 molecules, ranked by XP GlideScore, were purchased from commercial vendors for experimental validation.

### QUANTIFICATION AND STATISTICAL ANALYSIS

#### Statistical analysis

The data was analyzed using GraphPad Prism or R (version 4.3.1) unless otherwise specified. The means ± standard errors are presented in the results. Student t-tests were used to identify differences between the two groups and Pearson correlation comparing two conditions as indicated. For multiple groups, one-way analysis of variance (ANOVA) tests was used. Statistical analyses were based on data collected from at least three independent experiments. P<0.05 indicated statistically significant differences.

#### RNA-seq data processing

RNA-seq data processing was performed using Nextflow nf-core rnaseq pipeline version 3.6 ^58^. The overall quality of sequencing reads was evaluated using FastQC (v.0.11.9) and Trim Galore, a wrapper tool around Cutadapt and FastQC. Sequence alignments to the reference human genome (GRCh38) were performed using STAR and the gene expression was quantified using Salmon with the comprehensive annotations made available by Gencode (v29 GTF File). Samples were adjusted for library size and normalized with the variance stabilizing transformation (vst) in the R statistical environment using DESeq2 (v1.40.2) package ^59^. When performing differential expression analysis between groups, we applied the embedded independent filtering procedure to exclude genes that were not expressed at appreciable levels at least in three samples considered. P values were corrected for multiple testing using the Benjamini-Hochberg false discovery rate (FDR) procedure, with the significance threshold set to 0.05. If not otherwise specified, all GSEAs were performed using clusterProfiler R Package ^60^. Gene-set collections were retrieved from the Molecular Signature Database (MSigDB) ^61^.

#### LC-MS/MS data analysis

DIA raw data from proteomes were searched with DIA-NN version 1.8.1 with default settings against a deep learning-based predicted library generated from the human Uniprot database ^62^. For library generation, the FASTA sequences were digested with Trypsin/P with 1 missed cleavage, enabling N-terminal methionine excision and cysteine carbamidomethylation as a fixed modification. Precursor and fragment mass tolerance were determined automatically for each run separately and ranged from 10 to 20 ppm. The ‘report.pg_matrix.tsv’ table was used for further analysis.

BioID DDA raw files were processed with the MaxQuant software v.1.6.7.0 ^46^. Searches were performed against the Human UniProt database (June 2019) and a common contaminants database with a false discovery rate of 1% at both peptide and protein levels. Enzyme specificity was set as “Trypsin/P” with a maximum of 2 missed cleavages and 7 as the minimum length required for peptide identification. N-terminal protein acetylation, methionine oxidation, and lysine biotinylation were set as variable modifications, and cysteine carbamidomethylation as a fixed modification. Match between runs was enabled to transfer identifications based on mass and normalized retention times, with a matching time window of 0.7 min and an alignment time window of 20 min. Label-free protein quantification (LFQ) was performed with the MaxLFQ algorithm where a minimum peptide ratio count of 1 was required for quantification ^63^.

All the resulting datasets were analyzed with the Perseus software v.1.6.2.3 ^47^. Data were pre-processed by removing proteins only identified by site, reverse hits, and potential contaminants, and log2 transformation of LFQ intensities was applied. Biological triplicates of each experimental condition were grouped, and proteins were filtered for a minimum of 3 valid values in at least one group. Missing data points were replaced by imputation from a normal distribution with 0.3 widths and 1.8 downshift, and a two-sided two-sample Student’s t-test (permutation-based FDR, 250 randomizations) was used to identify significant changes in protein intensity between the experimental conditions.

#### ChIP-seq data processing and analysis

The genomic DNAs were processed using the NebNext Ultra II DNA library preparation kit with sample purification beads and sequenced on Illumina NextSeq 2000 with single-end, 120-base-pair long reads. The ChiP-seq reads were processed using the Nextflow nf-core open-source pipeline chipseq (version 1.2.2) ^64^ for quality control, alignment and peak-calling. The overall quality of the FASTQ files was first evaluated by FastQC (v.0.11.9) and the reads were mapped to the reference human genome (GRCh38) using bwa (version 0.7.17) software ^65^. From the alignment files narrow or broad enriched ChIP regions were evaluated using MACS2 ^66^ software and created corresponding normalised bigWig files scaled to 1 million mapped reads using BEDTools ^66^.

To compare the peak scores between FOXA1 and FOXA2 chip-seqs a merged peak set was created using BEDTools ^43^ merge function, and the signal scores 1000bp or 3000bp upstream and downstream of the identified peaks were computed using UCSC tool bigWigAverageOverBed ^67^. To compare the peak intensities between different conditions (Eg: KD vs parental or T-compound vs DMSO) computeMatrix tool from python deepTools ^44^ was used and the scores were then visualised as heatmaps and density plots using plotHeatmap and plotProfile tools respectively.

To predict the enrichment of known transcription factor motifs in the identified peaks, a list of cell type-specific peak coordinates of FOXA1 and FOXA2 binding regions was generated, and the genomic regions were scanned by HOMER (v5.0.1) software findMotifsGenome function ^45^. Motif enrichment analyses were centered on the 200 bp surrounding the peak summit. Chi-square statistical test was used to show the significant relative enrichment of transcription factor motifs in T-compound treated cells relative to control treatment.

---

**Figure S1.**
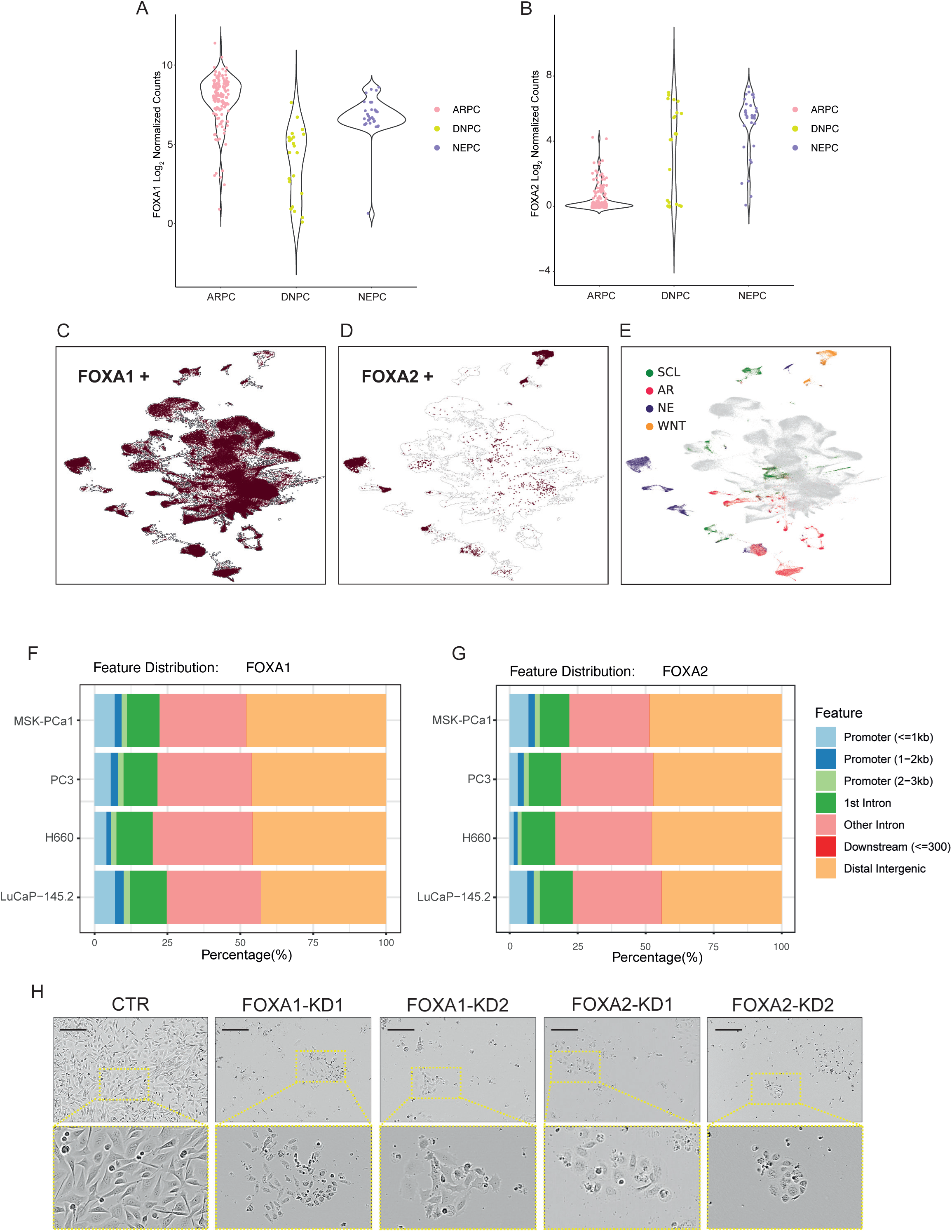
FOXA1 and FOXA2 expression in human prostate cancer RNAseq data and chromatin occupancy in prostate cancer models, related to Figure 1. (**A, B**) Violin plots indicate FOXA1 (A) and FOXA2 (B) gene expression levels in bulk RNA sequencing atlas. (**C-E**) UMAP representations of single-cell prostate cancer atlas highlights the expression of FOXA1 and FOXA2 and the different clusters corresponding to the castration-resistant prostate cancer subtypes, i.e., AR-driven prostate cancer (AR), WNT, stem-cell like (SCL), and neuroendocrine prostate cancer (NEPC). (**F, G**) Distribution of FOXA1 and FOXA2 peaks across the genome in the indicated cell line models. (**H**) Representative pictures using phase contrast microscopy of PC3 control cells (CTR) and cells with knockdown (KD) of either FOXA1 or FOXA2 using two different hairpin RNAs (bar represents 50µm).

**Figure S2.**
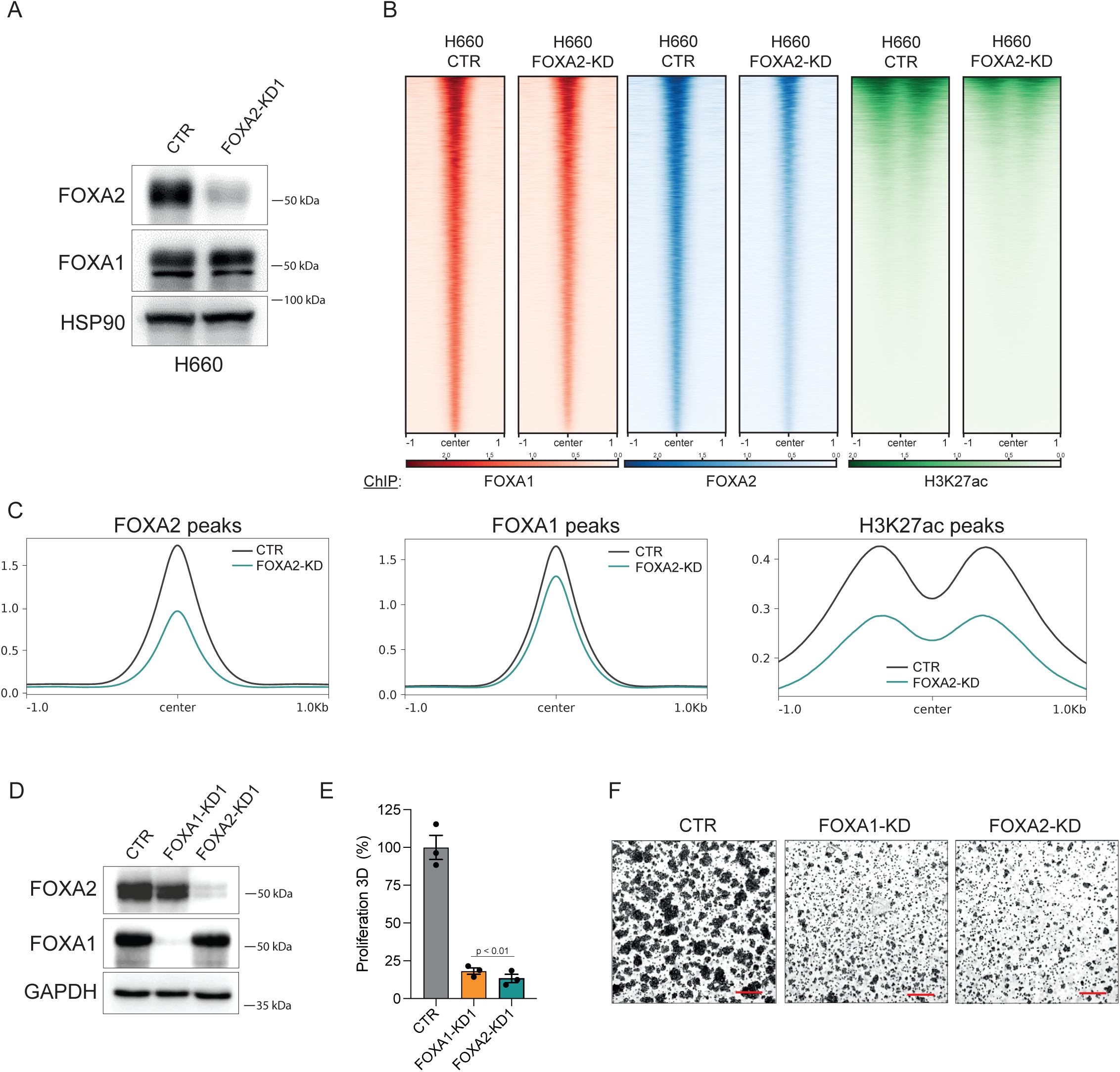
Knockdown of FOXA2 reduces FOXA1 chromatin occupancy and H3K27 acetylation (H3K27ac), as well as cell growth in neuroendocrine H660 cells, related to Figure 1. (**A**) Immunoblot of indicated proteins upon FOXA2 knockdown (KD) in H660 cells. (**B**) ChIPseq heatmaps for FOXA1, FOXA2, and H3K27ac of H660 control cells or knockdown (KD) of FOXA2 as indicated. (**C**) Corresponding change in FOXA1, FOXA2, and H3K27ac global profiles. (**D**) Immunoblot of indicated proteins upon FOXA2 and FOXA1 knockdown (KD) in H660 cells. (**E, F**) Quantification of clonogenic assay and corresponding representative pictures of H660 control cells and knockdown of FOXA2 and FOXA1 (bar represents 100µm). P-value 0.002 (student’s t-test). See also Figure S9B.

**Figure S3.**
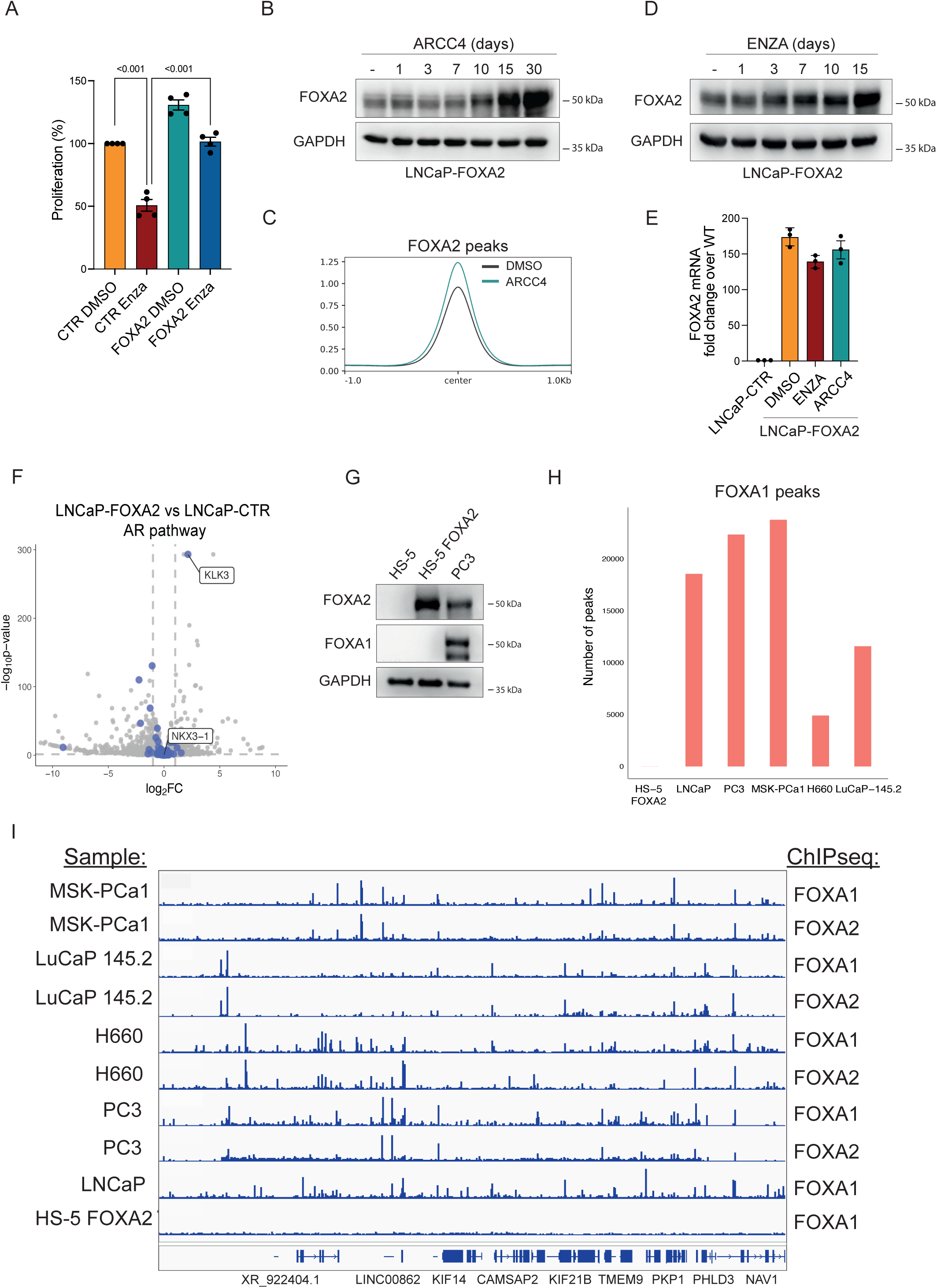
FOXA2 over-expression in LNCaP cells promotes AR-independent growth, related to Figure 2. (**A**) Proliferation assay showing growth reduction in LNCaP cells upon ENZA 10µM treatment in the presence (FOXA2) and absence of FOXA2 (CTR). P-values <0.001 for both ENZA vs DMSO and ENZA CTR vs ENZA FOXA2 (student’s t-test). (**B**) Immunoblot for the indicated proteins of LNCaP cells over-expressing FOXA2 (LNCaP-FOXA2) exposed to ARCC4 1µM for the indicated number of days. (**C**) Corresponding analysis of global chromatin binding in LNCaP-FOXA2 cells with and without ARCC4 1µM treatment for 30 days. (**D**) Immunoblot for the indicated proteins of LNCaP cells over-expressing FOXA2 (LNCaP-FOXA2) exposed to ENZA 10µM for the indicated number of days. (**E**) mRNA expression of FOXA2 by quantitative (q)PCR in LNCaP-CTR and in cells over-expressing FOXA2 with or without ENZA 10µM and ARCC4 1µM treatment for 30 days. (**F**) Changes of AR target proteins in response to FOXA2 over-expression in LNCaP cells. (**G**) Immunoblot for the indicated proteins and cell lines. HS-5 represents a fibroblast cell line lacking FOXA1/2 expression. (**H**) The number of peaks for FOXA1 was measured across the indicated cell lines and organoid models in ChIPseq data using one replicate. (**I**) Exemplified track across different models highlighting the lack of peaks for FOXA1 in HS-5 over-expressing FOXA2.

**Figure S4.**
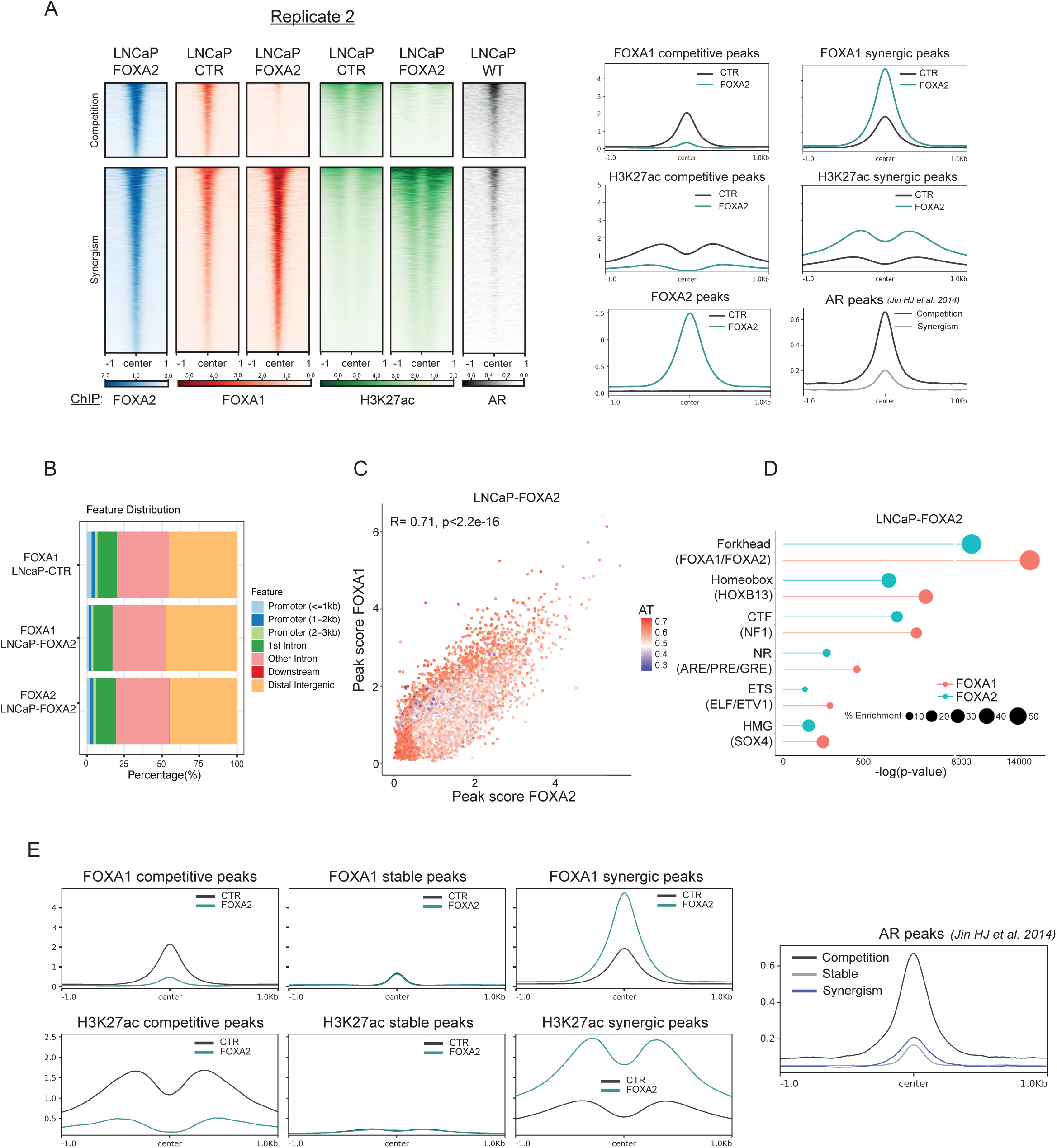
FOXA2 genomic localization in FOXA2 over-expressed LNCaP cells, related to Figure 2. (**A**) Heatmaps and global ChIPseq profiles for FOXA1, FOXA2, and H3K27ac of the second replicate corresponding to the experiment in Figure 2D. (**B**) Distribution of peaks across the genome related to FOXA1 and FOXA2 in LNCaP control (CTR) cells and cells over-expressing FOXA2 (LNCaP-FOXA2). (**C**) Scatter plot comparing peak scores for FOXA1 and FOXA2 in LNCaP cells over-expressing FOXA2 (LNCaP-FOXA2) in duplicates. AT content of the peaks is indicated by color-code. Pearson correlation coefficient and corresponding p-values are indicated for genes with significant changes. (**D**) Corresponding lollipop plot highlighting the enrichment of selected DNA binding motifs (upper part) and corresponding transcription factors (indicated in brackets, lower part) in FOXA1 and FOXA2 ChIPseq data (see also table S1_1). (**E**) Three-tiered distribution of FOXA1 chromatin occupancy changes in control LNCaP cells and cells over-expressing FOXA2. Corresponding changes in H3K27ac and association with previously published AR binding sites. See also Figures 2D & S4A.

**Figure S5.**
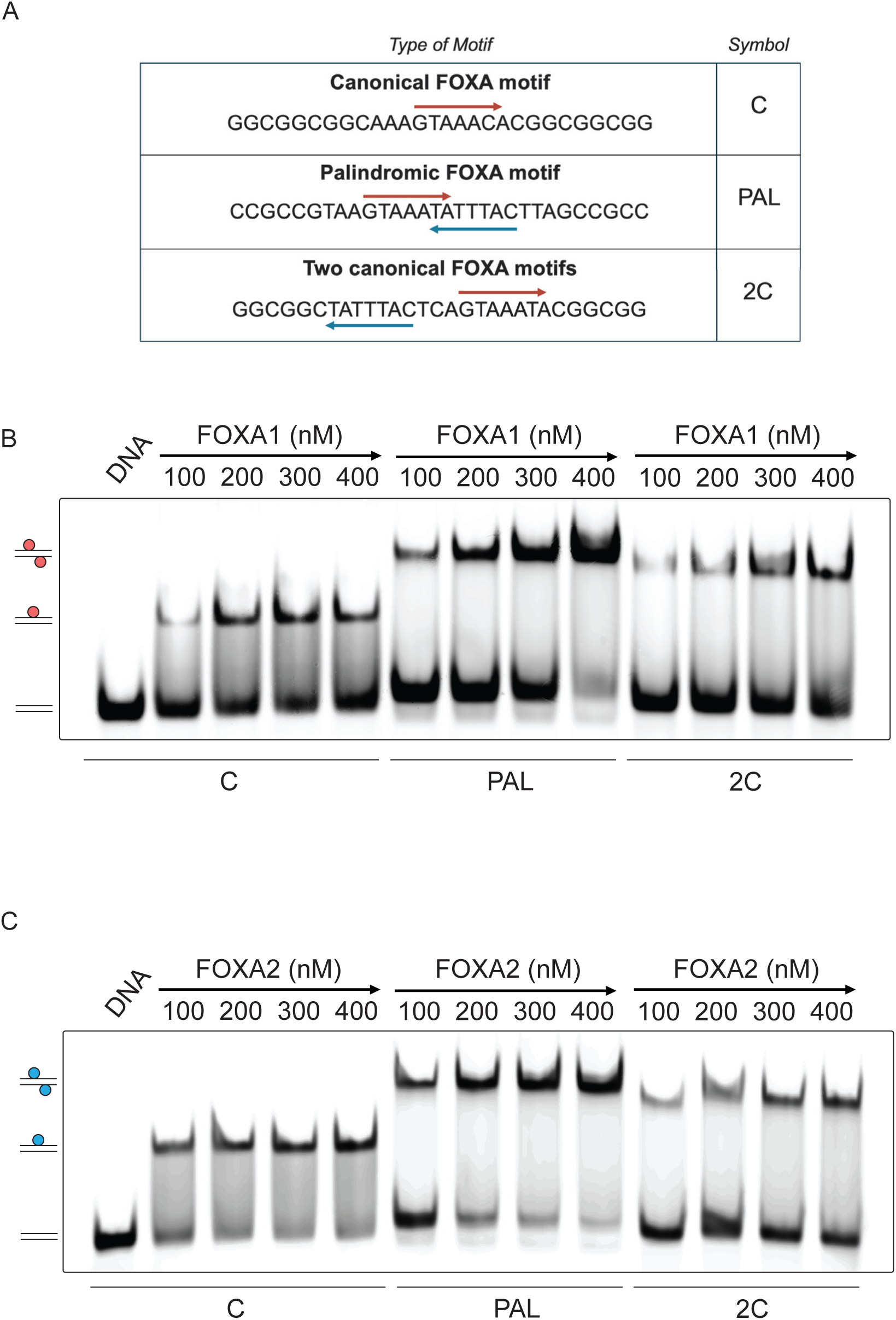
Interaction of the recombinant FOXA1 and FOXA2 forkhead domains with DNA oligonucleotides, related to Figure 2. (**A**) Outline of the DNA nucleotides harboring different FOXA motifs: the canonical motif (C), extended palindromic motif (PAL), and the two canonical motifs separated by three nucleotides (2C). The latter two are known to enable homodimerization of FOXA1 forkhead domains. (**B**) Electrophoretic Mobility Shift Assay (EMSA) with oligonucleotides harboring either the canonical motif (C), the extended palindromic motif (PAL), and the two canonical motifs separated by three nucleotides (2C) with increasing concentrations of the FOXA1 forkhead domain as indicated. (**C**) Corresponding assays using the FOXA2 forkhead domain.

**Figure S6.**
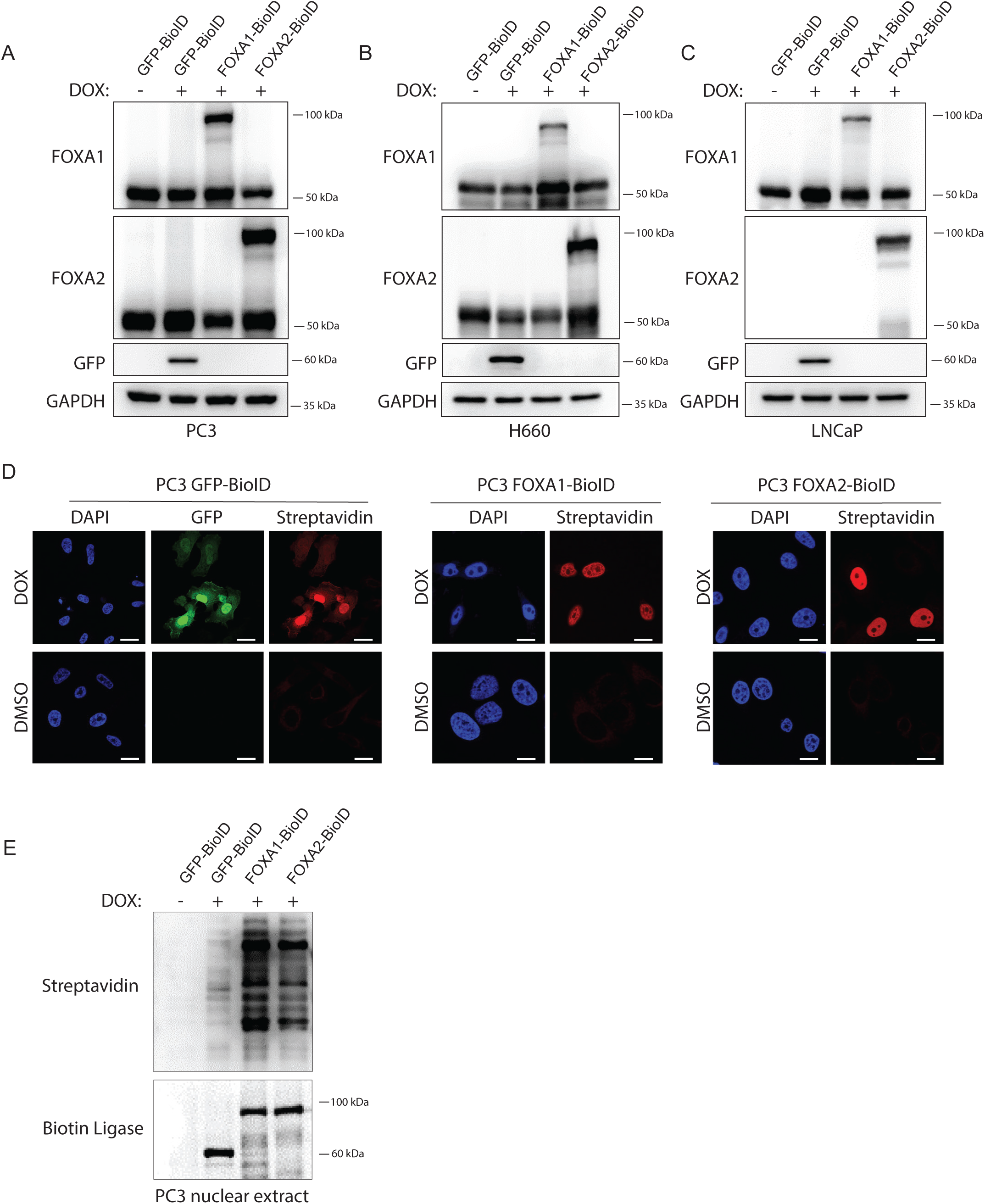
Characterization of cell lines used for BioID, related to Figure 3. (**A-C**) Immunoblot analysis of the indicated protein in doxycycline (DOX)-inducible GFP-control, FOXA1, or FOXA2-tagged biotin ligase (BioID) in the indicated cell lines. (**D**) Corresponding immunofluorescence for biotinylated proteins upon induction with and without DOX in PC3 cells over-expressing GFP-control, FOXA1, or FOXA2-tagged biotin ligase (BioID). (**E**) Immunoblot analysis for biotinylated proteins and biotin ligase in PC3 cells expressing GFP-, FOXA1-, or FOXA2-tagged biotin ligase (BioID).

**Figure S7.**
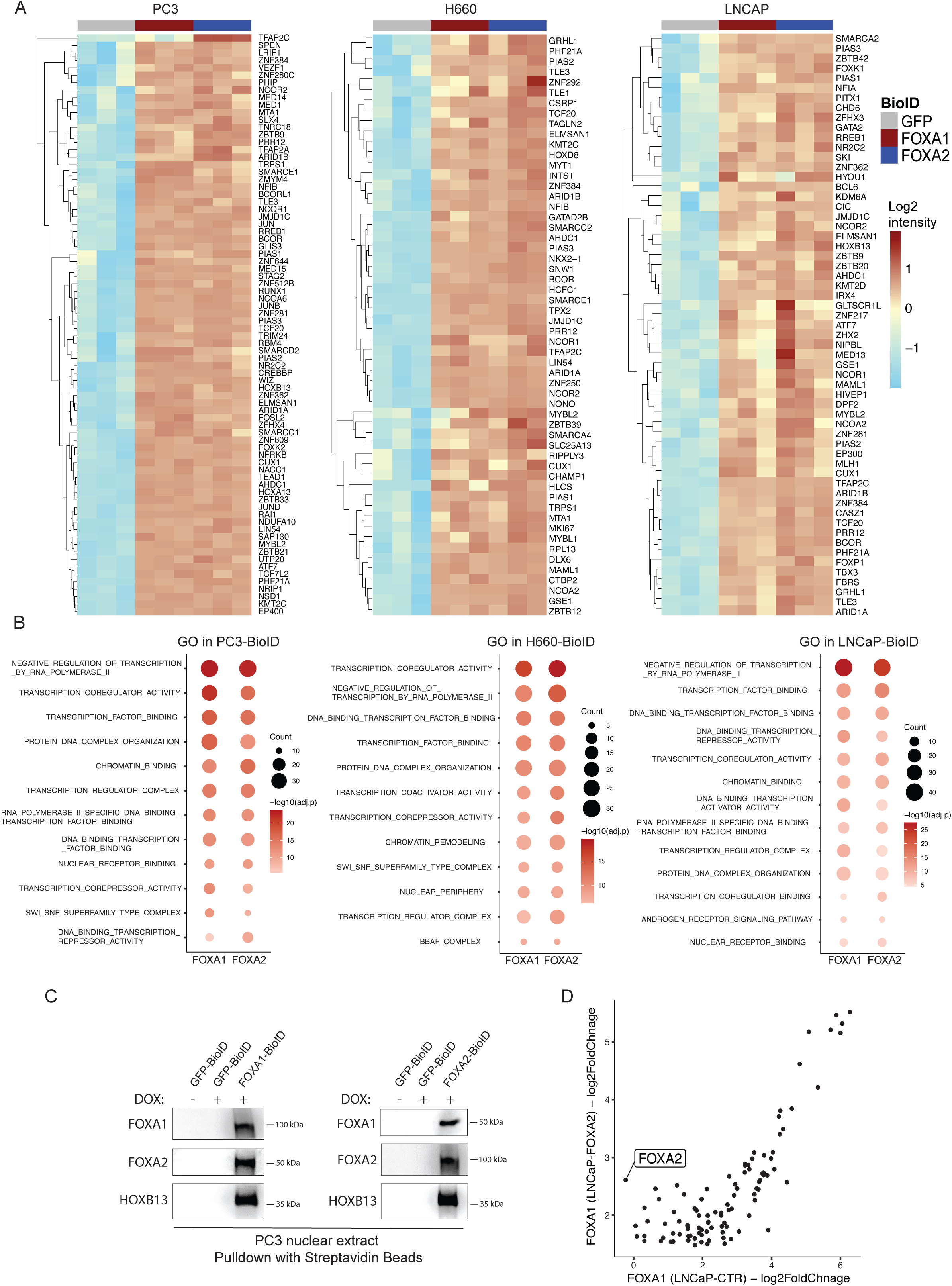
Quality checks on BioID data in the different cell lines, related to Figure 3. (**A**) Heatmaps of un-normalized log2 protein enrichment values in the indicated BioID data using GFP, FOXA1, FOXA2 data as a bait in the indicated cell lines. (**B**) Gene ontology analysis highlighting the top 10 enriched signatures of FOXA1 and FOXA2 BioID data as compared to GFP control in the indicated cell lines. (**C**) Immunoblot analysis for the indicated proteins after biotin-streptavidin pulldown in PC3 cells expressing either GFP-, FOXA1-, or FOXA2-tagged biotin ligase. (**D**) Scatter plot comparing enrichment scores for FOXA1 BioID targets over GFP control in LNCaP control (CTR) and LNCaP cells over-expressing FOXA2 (LNCaP-FOXA2). Pearson correlation coefficient and corresponding p-values are indicated.

**Figure S8.**
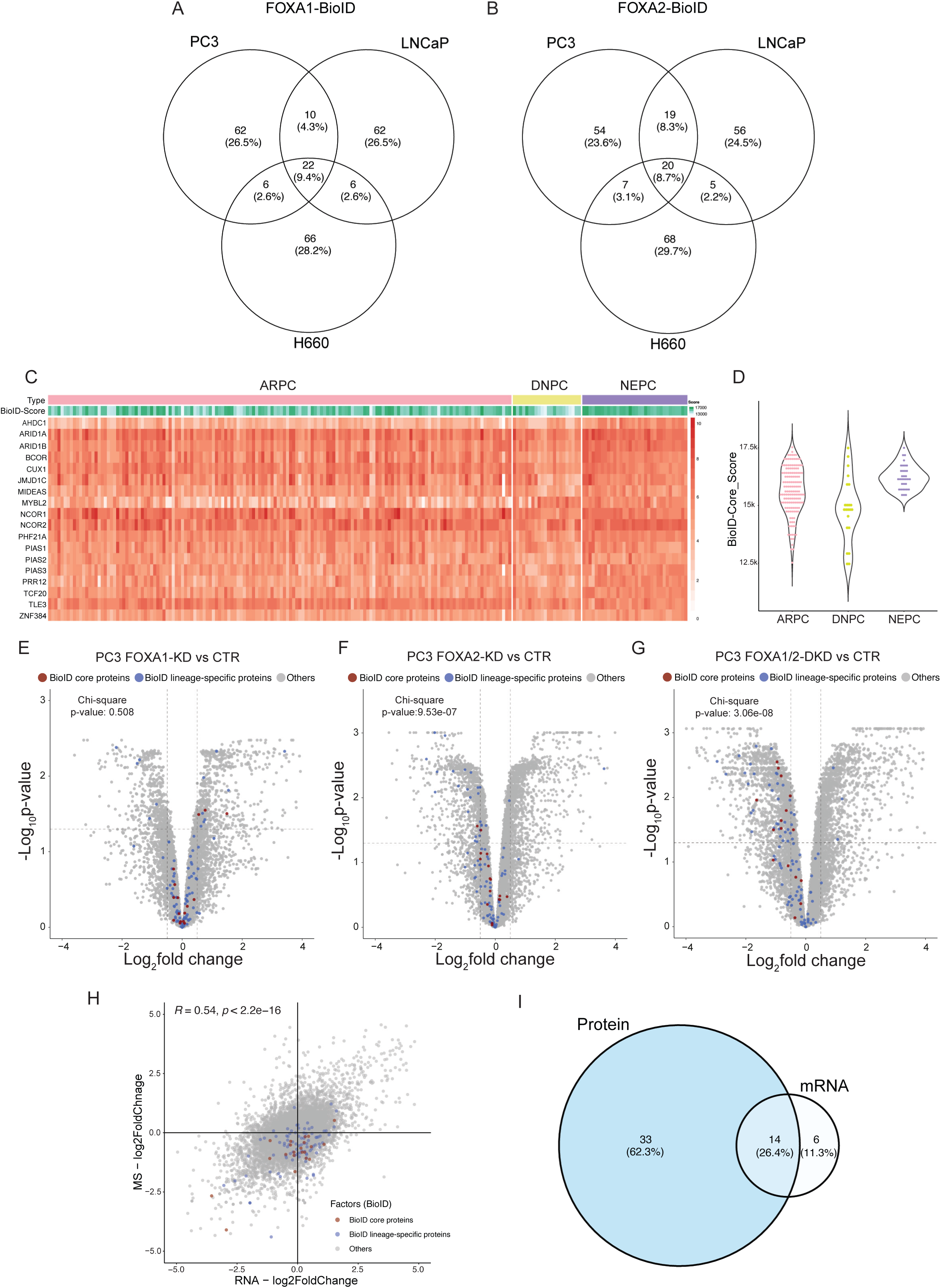
Characterization of BioID interactors, related to Figure 3. (**A**, **B**) Venn diagrams of the top 100 BioID-enriched proteins for FOXA1 (**A**) and FOXA2 (**B**) in the indicated cell lines. (**C**) Heatmap of the gene expression of common BioD interactors (BioID Core) across castration-resistant prostate cancer (ARPC), neuroendocrine prostate cancer (NEPC), and double-negative prostate cancer (DNPC) (see also Figure 1A). (**D**) Corresponding gene set enrichment analysis of the BioID Core signature. (**E-G**) Volcano plot of mass-spectrometry data of PC3 cells comparing FOXA1 knockdown (KD) with control (CTR) (**E**), FOXA2 KD with CTR (**F**), and FOXA1/2 double knockdown (DKD) with CTR (**G**). The top 100 BioID-enriched proteins are labeled, and the significance of enrichment in downregulated proteins is indicated by Chi-square analysis and corresponding p-value. (**H**) Scatter plot comparing protein and mRNA expression changes in PC3 in FOXA1/2 DKD versus CTR. The top 100 BioID-enriched proteins are highlighted. Pearson correlation coefficient of all data points and corresponding p-values are indicated. (**I**) The corresponding Venn diagram overlaps the top 50 BioID-enriched proteins regulated at the mRNA and protein level in PC3.

**Figure S9.**
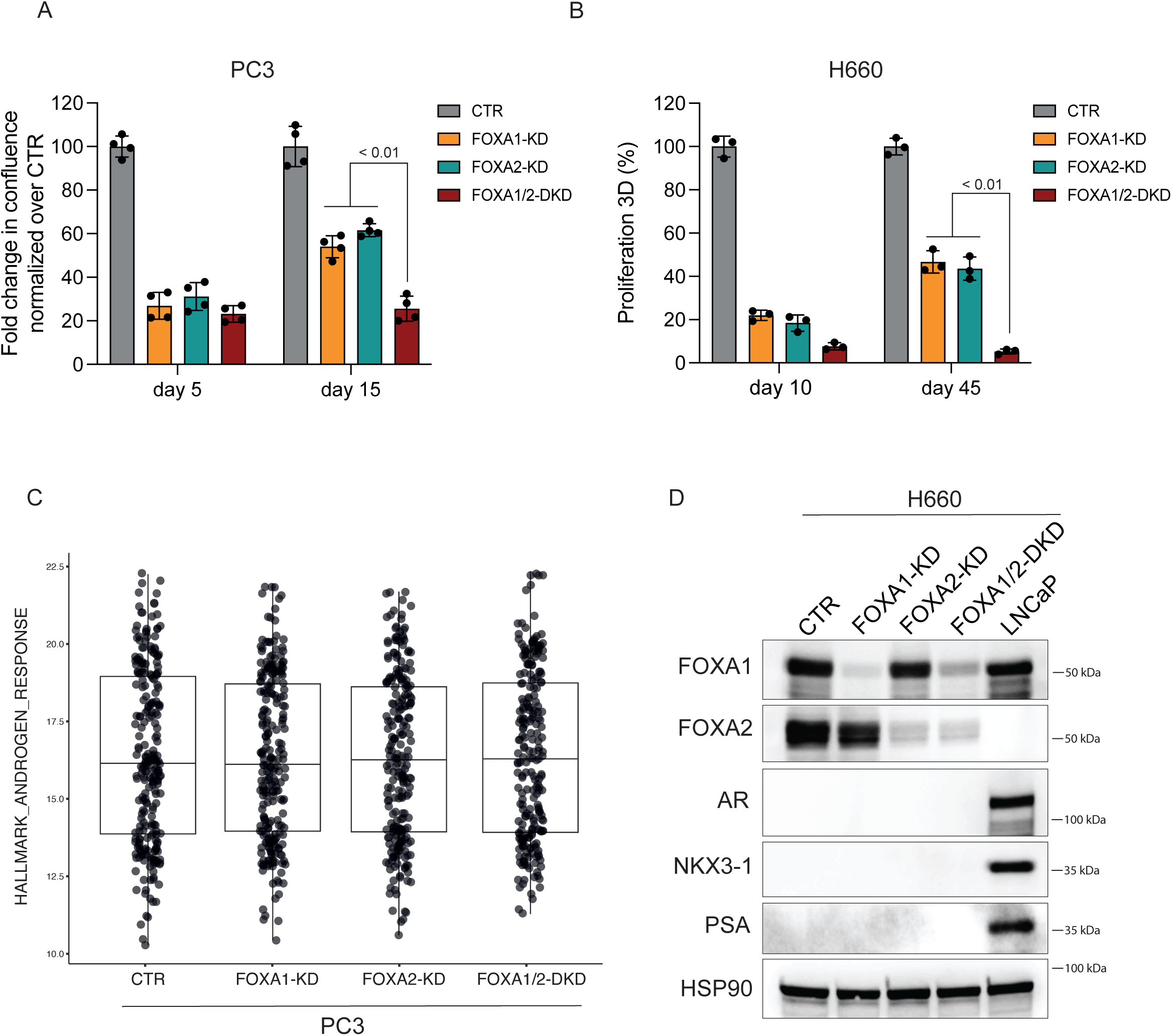
Effect of FOXA1 and FOXA2 single- and double-knockdown on cell proliferation and AR signaling in PC3 and H660 cells, related to Figure 3. (**A**) Proliferation assay over 5 and 15 days of PC3 cells of control (CTR) cells, FOXA1 and FOXA2 single knockdown (KD) and double knockdown (DKD) using one short hairpin RNA for each gene. (**B**) Corresponding proliferation assay over 10 or 45 days of H660 cells. (**C**) Protein abundance changes of AR and downstream targets in the mass-spectrometry data related to CTR, FOXA1 and FOXA2 single KD and DKD in PC3 cells. See also Figure 3D-F. (**D**) Corresponding immunoblot analysis for AR and the downstream targets NKX3-1 and PSA in H660 control (CTR) cells, FOXA1 and FOXA2 single knockdown (KD) and double knockdown (DKD) using one short hairpin RNA for each gene. LNCaP cells are included as a positive control.

**Figure S10.**
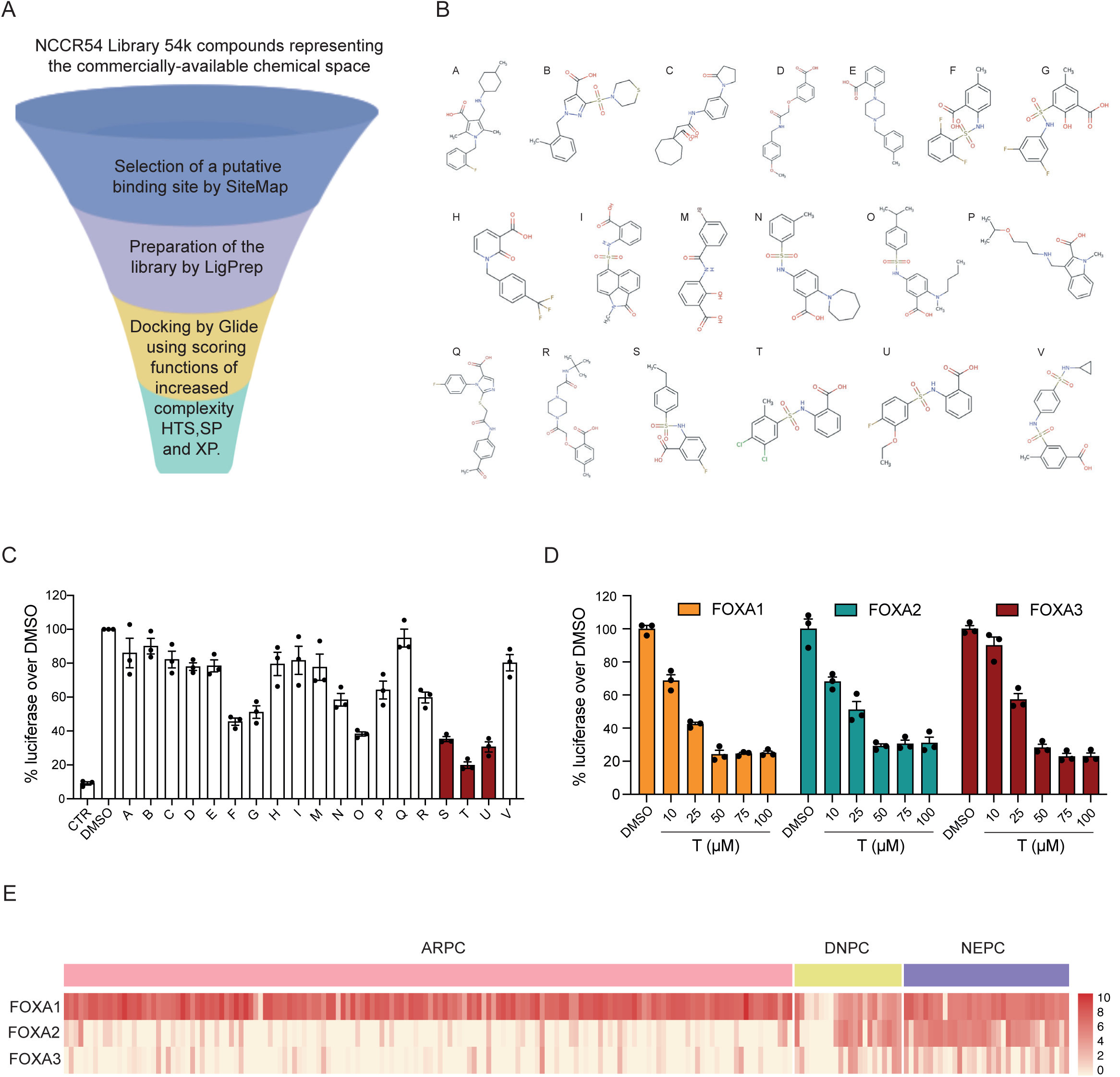
Small molecules emerging from virtual drug screening, related to Figure 4. (**A**) Graphical representation of the virtual screening protocol applied (see Figure 4A). (**B**) Chemical structure of small molecules A-V emerging from virtual drug screening tested by the luciferase assay. (**C**) Luciferase reporter assay indicating the reduction in FOXA1-induced luciferase activity by the indicated small molecules tested at 100µM for 24 hours. (**D**) Luciferase reporter assay indicating the reduction in FOXA1, −2, or −3 induced luciferase activity by T compound at 100µM for 24 hours. (**E**) FOXA1, −2, and −3 mRNA expression in bulk RNA sequencing atlas data of AR-positive castration-resistant prostate cancer (ARPC), neuroendocrine prostate cancer (NEPC), and double-negative prostate cancer (DNPC), www.prostatecanceratlas.org ^23^.

**Figure S11.**
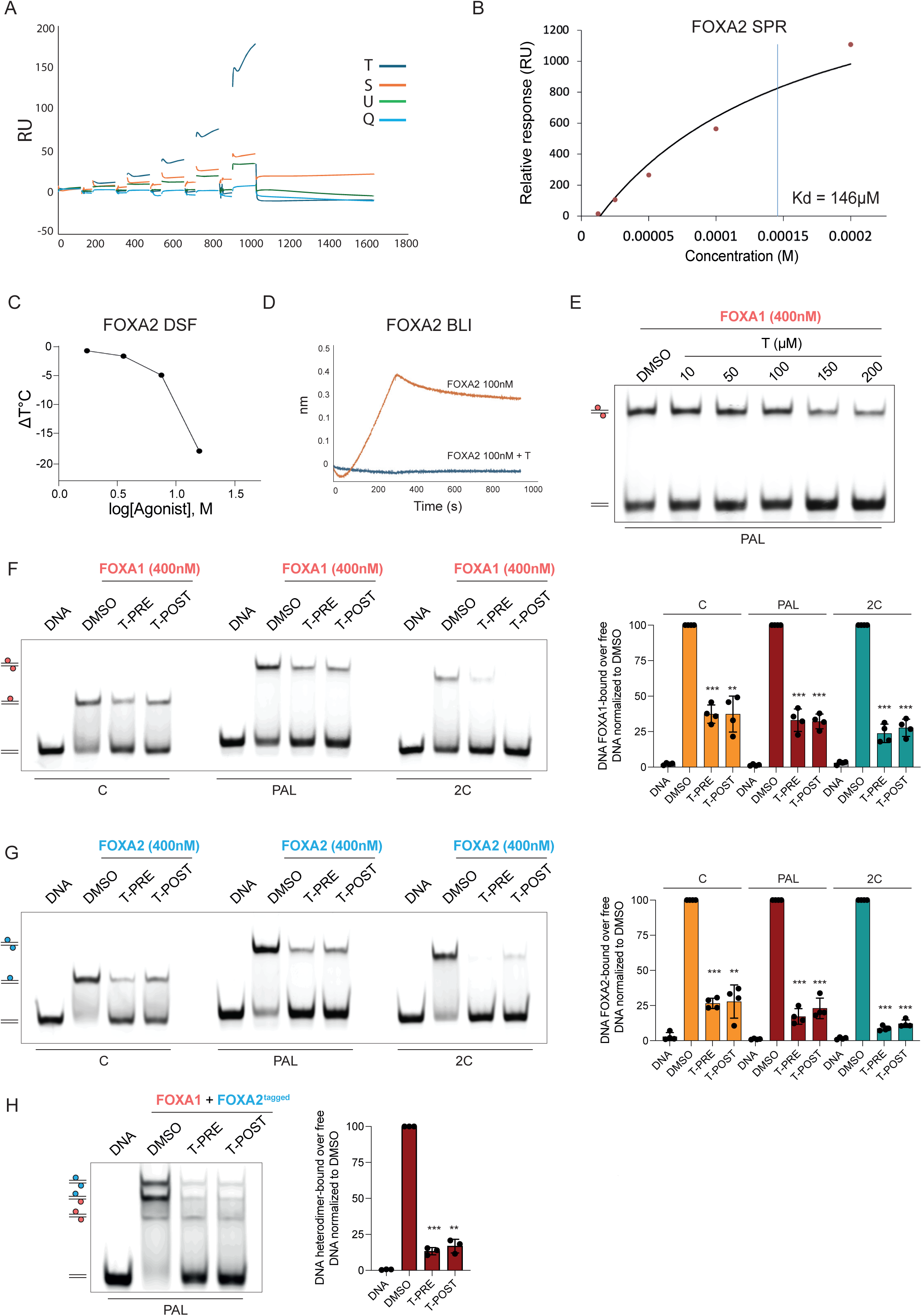
Effects of compound T on the forkhead domains of FOXA1 and FOXA2 using biophysical assays, related to Figure 4. (**A**) Sensograms for compounds T, S, U, and Q and the forkhead domain of FOXA1 in Surface Plasmon Resonance (SPR) experiments (see Figure 4 C). (**B**) SPR experiment showing the binding for the T compound using the immobilized recombinant forkhead domain of FOXA2. The dissociation constant (KD) is calculated by a steady state model and resulted to be 146µM. (**C**) Influence of different concentrations of T compound on the melting temperature of the recombinant FOXA2 forkhead domain measured by Differential Scanning Fluorimetry (DSF). (**D**) Ability of the FOXA2 forkhead domain to bind an immobilized double-strand DNA sequence in the presence and absence of 100µM of compound T by Bio-Layer Interferometry BLI. (**E**) Electrophoretic Mobility Shift Assay (EMSA) using the palindromic nucleotide DNA template and the FOXA1 forkhead domain with pre-incubation of T compound at the indicated concentration before adding the nucleotide template. (**F, G**) Corresponding assay using either the FOXA1 or FOXA2 forkhead domain, respectively, with either pre-incubation (PRE) of T as in (E) or the addition of T 200µM one hour after incubation of the forkhead domain with the DNA oligonucleotide (POST) as indicated. Quantification of the reduction of DNA-protein complexes induced by T 200µM in four replicates. Statistical significance is indicated by asterisks: *p < 0.05, **p < 0.01, ***p < 0.001. (**H**) Effect of T compound 200µM on heterodimer and corresponding quantification, see above and Figure 2F. Statistical significance is indicated by asterisks: *p < 0.05, **p < 0.01, ***p < 0.001.

**Figure S12.**
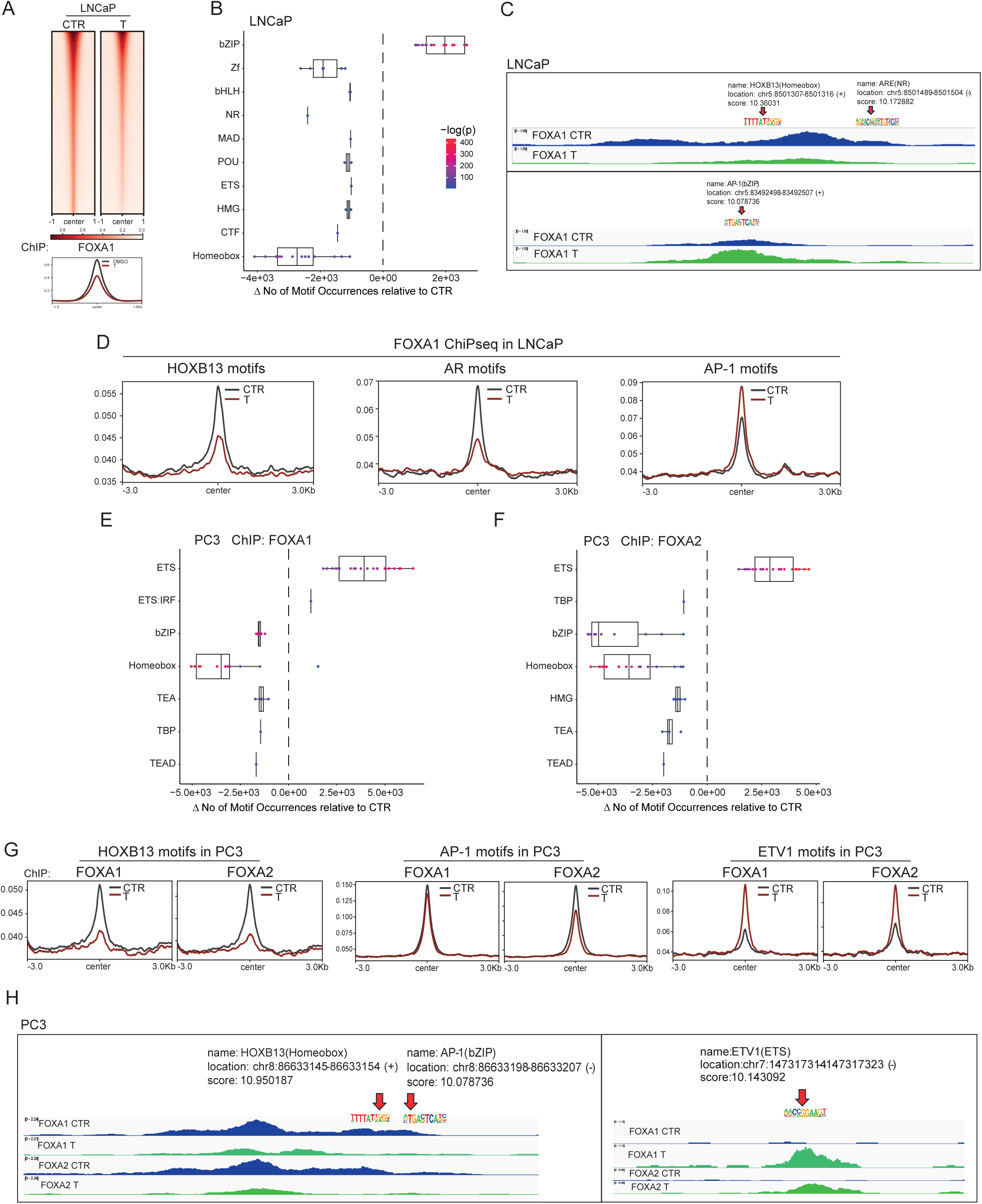
Effects of T compound on FOXA1 and FOXA2 genomic localization, related to Figure 4. (**A**) ChIPseq heatmaps and global profiles for FOXA1 in LNCaP cells treated with 100µM T compound for one day compared to control (DMSO, CTR). (**B**) Corresponding analysis of T compound’s depleted and enriched DNA motifs compared to control. (**C**) Exemplified changes of depleted and enriched DNA motifs as indicated. (**D**) Corresponding ChIPseq profiles of LNCaP cells treated with 100µM T and control (CTR) stratified according to the indicated DNA binding motifs. (**E, F**) Analysis of depleted and enriched DNA motifs in FOXA1 and FOXA2 ChIPseq data of PC3 cells treated with 100µM T compound for one day compared to control (see Figure 4H). (**G**) Corresponding ChIPseq profiles of PC3 cells treated with 100µM T and control stratified according to the indicated DNA binding motifs. (**H**) Exemplified changes of depleted and enriched DNA motifs as indicated.

**Figure S13.**
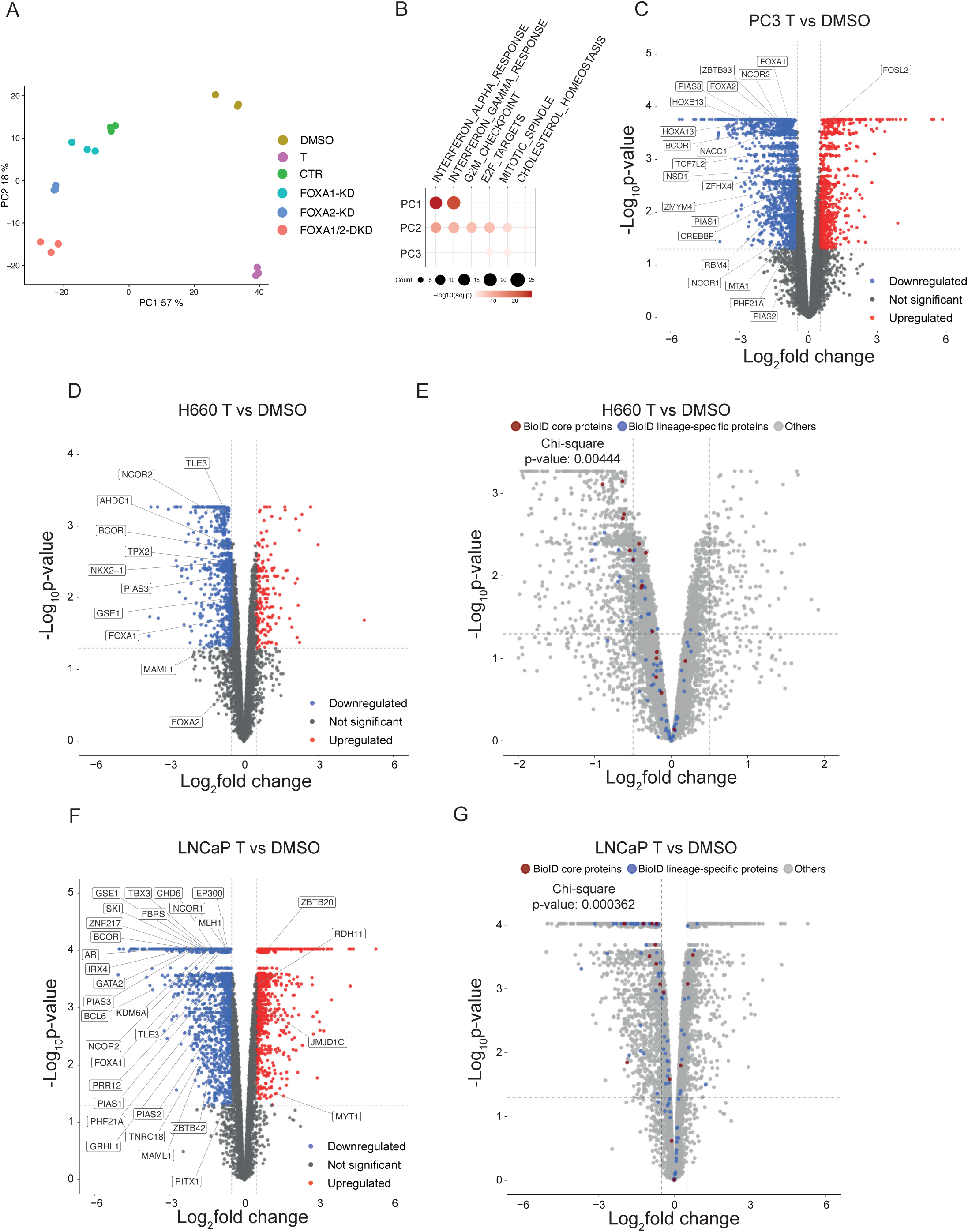
Effects of T compound on the proteome of PC3, H660 and LNCaP cells, related to Figure 4. (**A**) Principal component analysis of protein changes of FOXA1 and FOXA2 single (KD) and double knockdown (DKD) compared to PC3 cells treated with 100µM compound T or control DMSO for 2 days. (**B**) Corresponding gene set enrichment analysis of principal components (PC) 1-3. (**C**) The volcano plot of protein abundance changes in PC3 cells after incubation with 100 µM compound T for 2 days. Significant changes are labeled in the top 50 BioID interactors. (**D)** The volcano plot of protein abundance changes in H660 cells after incubation with 100µM compound T for 2 days. Significant changes in the top 50 BioID interactors are labeled. (**E**) The corresponding plot highlights the distribution of the top 100 BioID interactors (blue: lineage-specific interactors, red: core components). The significance of enrichment in downregulated proteins is indicated by Chi-square analysis and the corresponding p-value. (**F**) The volcano plot of protein abundance changes in LNCaP cells after incubation with 100µM compound T for 2 days. Significant changes in the top 50 BioID interactors are labeled. (**G**) The corresponding plot highlights the distribution of the top 100 BioID interactors (blue: lineage-specific interactors, red: core components). The significance of enrichment in downregulated proteins is indicated by Chi-square analysis and the corresponding p-value.

**Figure S14.**
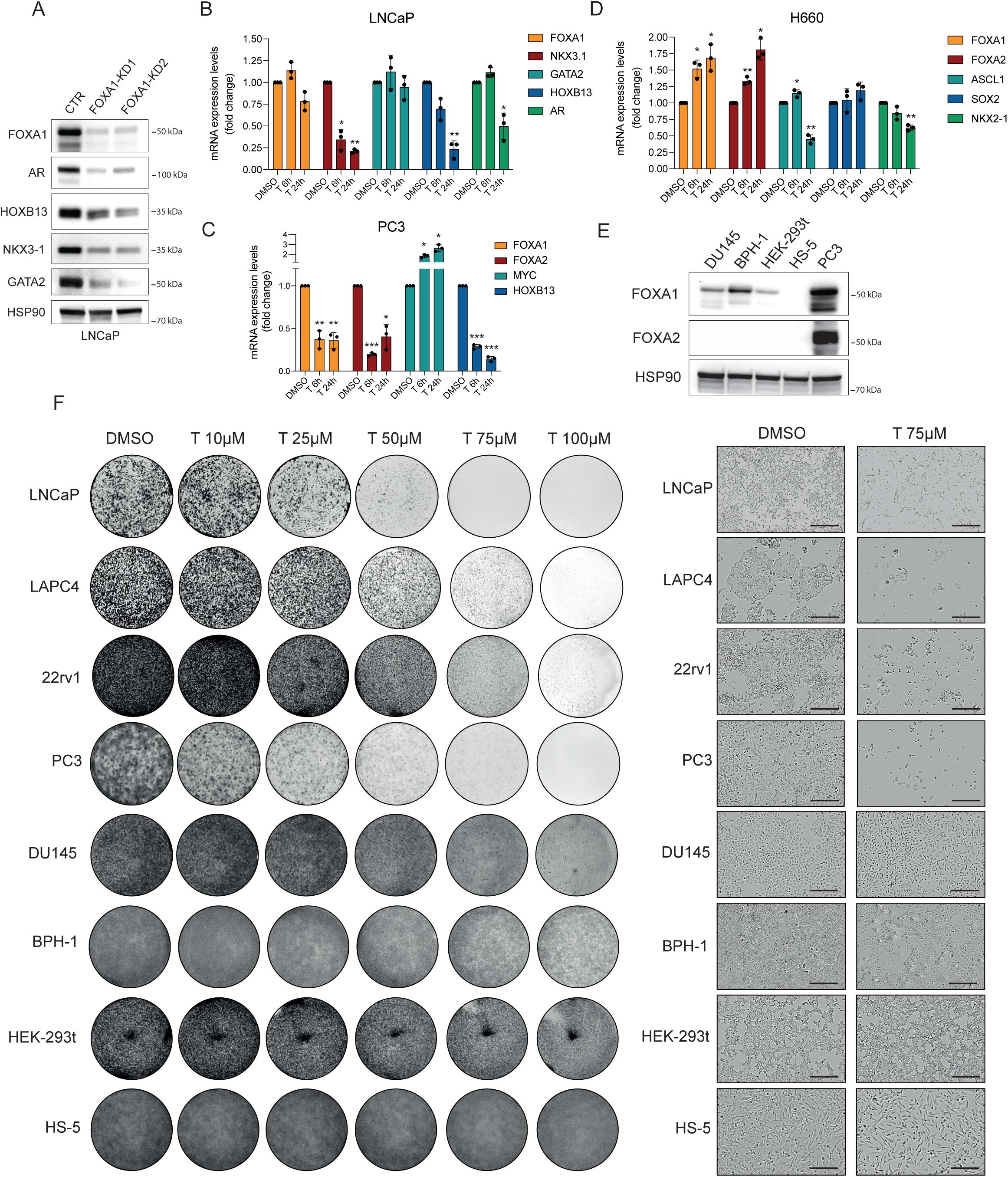
Transcriptional regulation of lineage-specific transcription factors and different sensitivity of cells to T compound, related to Figure 4. (**A**) Immunoblot analysis of LNCaP cells with knockdown (KD) of FOXA1 with two short hairpin RNAs probed for the indicated proteins. (**B, C, D**) Quantification of mRNA expression changes of the indicated transcription factors related to 100µM T compound treatment for 6 and 24 hours (h) by qPCR, see also Figure 4M-O for corresponding protein expression changes. (**E**) Immunoblot probed for the indicated proteins in the indicated cell lines with no or low FOXA1 or FOXA2 protein levels. PC3 cells serve as a reference for abundant FOXA1 and FOXA2 protein expression, see also Figure 1B. (**F**) A representative overview of cell culture wells related to the clonogenic assays of the indicated cell lines (left) with microscopic details (right) (see also Figure 4P). Bars represent 50µm.

**Table S1.** List of supplementary tables.

|  |  |
| --- | --- |
| table S1_1 | List of transcription factor motifs enriched (HOMER) in FOXA1 and FOXA2 Chip-seq in different cell lines (PC3, H660, MSK-PCa1, LuCaP- 145.2, LNCaP and VCAP) |
| table S1_2 | GSEA of Hallmark Biological pathways in CRPCs (FOXA2<sup>high</sup> vs FOXA2<sup>low</sup>) |
| table S2 | GSEA of Hallmark Biological pathways in LNCaP cells (Parental, ENZA and ARCC4) |
| table S3_1 | FOXA1 and FOXA2 BioID (PC3, H660 and LNCaP) |
| table S3_2 | GSEA of GO Biological pathways enriched with in core proteins |
| table S3_3 | Differential expression in PC3 cell line - RNA (DKD vs PLKO) and protein (DKD/shFOXA1/shFOXA2 vs PLKO) |
| table S4_1 | List of transcription factor motifs enriched (HOMER) with T compound treatment (PC3 and LNCaP) |
| table S4_2 | GSEA of Hallmark Biological pathways enriched in the first three principal components of DKD/shFOXA1/shFOXA2 and T compound treatment (PC3) |
| table S4_3 | Differential expression - T vs DMSO RNA (PC3) and Protein (PC3, H660 and LNCaP) |
| table S4_4 | GSEA of Hallmark Biological pathways enriched in different cell lines (PC3, H660 and LNCaP) treated with T compound |

